# Bellinger River Virus 2: The Emergence of a Second Novel Pathogenic Nidovirus in Bellinger River Snapping Turtles (*Myuchelys georgesi*)

**DOI:** 10.64898/2026.11.27.769811

**Authors:** Lauren Lim, Bethan J. Lang, Josephine Rieken, Kate Parrish, Jane Hall, Karrie Rose, Edward C. Holmes

## Abstract

In 2015 a major population decline in the critically endangered Bellinger River snapping turtle (*Myuchelys georgesi*; New South Wales, Australia) was attributed to the circulation of the serpentovirus Bellinger River virus (BRV). A decade later, we report the identification of a second novel serpentovirus in wild Bellinger River snapping turtles, herein referred to as Bellinger River virus 2 (BRV2), that is also associated with mortalities and further population impacts. Ocular and lacrimal gland lesions were consistent across disease events, although tissue tropism and distribution of internal gross and microscopic lesions varied between BRV and BRV2 infections. We employed a metatranscriptomic approach to identify and characterise BRV2. Phylogenetic analysis revealed that BRV and BRV2 form a distinct Bellinger River virus clade within the subfamily *Serpentovirinae* (family *Tobaniviridae*, order *Nidovirales*). However, given the extent of sequence divergence between the original BRV detected in 2015 and BRV2, which exhibit ∼70% nucleotide identity across the full viral genome, we propose that BRV2 did not emerge directly from BRV during the period of outbreak observation and instead arose from a recent cross-species transmission (spillover) event within the Bellinger River ecosystem. The emergence of BRV2 in *M. georgesi* suggests that Bellinger River viruses remain an ongoing threat to the conservation of the species and emphasises the need for continued effort in the biotic and abiotic surveillance of the river and its wildlife. Such surveillance may also aid in improving our understanding of the origins, transmission, and drivers of serpentovirus emergence within the river ecosystem.

**IMPACT STATEMENT:** The eponymously named Bellinger River snapping turtle (*Myuchelys georgesi*) is an endemic freshwater turtle that has been the subject of more than a decade of conservation efforts. While it was once considered an abundant species in the Bellinger River, the emergence of a novel nidovirus in 2015 resulted in a 90% population decline. Since this dramatic decline, population monitoring, breeding programs and pathogen surveillance for the aetiologic Bellinger River virus (BRV) have been in effect. In recent years, *M. georgesi* mortalities have been reported with pathologies similar to BRV; however, PCR diagnostic assays suggest the presence of an additional infectious agent. Herein, we characterised a second novel pathogenic Bellinger River virus 2 (BRV2) in turtles sampled from mortality events in 2022 and 2024 that likely emerged following an independent cross-species transmission event from a currently unknown reservoir. Together, findings contribute to the conservation of a critically endangered native Australian species.

**DATA SUMMARY:** The raw sequence reads generated here are available on the NCBI SRA under BioProject PRJNA1563736 and accession numbers SRR41835701-SRR41835716. All viral genomes and corresponding sequences assembled in this study have been deposited in GenBank under accession numbers QB391510-QB391512.

## INTRODUCTION

In 2015, the endangered and eponymously named Bellinger River snapping turtle (*Myuchelys georgesi*) suffered a 90% population decline over a course of six weeks due to the emergence and spread of a novel nidovirus denoted Bellinger River Virus (BRV) (1, 2). This catastrophic outbreak predominantly impacted sub-adult and adult *M. georgesi* turtles, without affecting other species within the Bellinger River ecosystem. The sole extant wild population of *M. georgesi* is estimated to comprise of only 100-200 individuals (3), and remains dependent on conservation efforts including ongoing population monitoring, community water quality surveillance (4), and captive breeding programs to facilitate population numbers and maintain population genetic diversity (5). Extended monitoring in the five years post-outbreak also revealed limited prevalence of BRV in wild *M. georgesi*, with no detections of the virus in the Bellinger River system since November of 2018.

BRV is a member of the order *Nidovirales* (family *Tobaniviridae*, subfamily *Serpentovirinae*; species *Pregotovirus myuchelyis*), and possesses a single-stranded, positive-sense RNA genome of approximately 31kb in length. BRV infection in *M. georgesi* is reported to present with bilateral conjunctiva inflammation, average to poor body condition, weakness and some animals displaying tan foci on the skin of the ventral thighs or hind limb paresis (2). While not all members of the *Tobaniviridae* are known to cause disease, the majority of the *Serpentovirinae* are associated with respiratory disease and pneumonia, including those in captive snakes and pythons, shingleback nidovirus 1, and veiled chameleon serpentovirus (6–8). In contrast, clear nasal discharge was the only reported respiratory manifestation in *M. georgesi* infected with BRV (9). Importantly, no other infectious agents have been reported to infect *M. georgesi*, while BRV has been detected in a small number of Macquarie River turtles (*Emydura macquarii*), another freshwater turtle species found along portions of the Bellinger River (5).

Population fluctuations have been reported in wild *M. georgesi* along the Bellinger River, coinciding with mortalities of turtles testing negative in BRV-specific PCR assays but positive in turtle serpentovirus RT-qPCR assays (Parrish *et al*. in preparation). These observations prompted further investigation into the potential viral aetiology of this emergent and uncharacterised disease in *M. georgesi*. In recent years, the increasing use of metatranscriptomics (i.e., total RNA sequencing) has offered a largely unbiased approach to the surveillance and discovery of known and novel infectious agents in wildlife (10, 11).

Using this approach, we identified and characterised a second novel Bellinger River virus in *M. georgesi*.

## METHODS

### Animal Ethics

Turtles collected for veterinary health and welfare assessments during the 2015 mortality event were euthanised based on their grave prognoses and subject to gross and histological post-mortem examination at the Australian Registry of Wildlife Health, Taronga Conservation Society Australia, Sydney. Two animal ethics committees were consulted and confirmed that research authorities were not required during active wildlife morbidity and mortality investigations. However, samples collected in follow-up surveillance were approved in Taronga Conservation Society Australia’s Animal Ethics Committee Research Authority 3e1015. Turtles found exhibiting clinical disease during 2022 and 2024 population surveys were collected under the NSW Department of Climate Change, Energy, the Environment and Water Animal Ethics Committee Animal Research Authority 180904/05. Samples were collected in accordance with the NSW National Parks and Wildlife Act 1974, section 132c, Scientific License numbers SL102155 and SL100104.

### Sample Collection

Tissue and swabs from 35 clinically diseased Bellinger River snapping turtles (*M. georgesi*) were collected across three years: 2015 (n=12), 2022 (n=5), 2024 (n=18). Samples collected in 2015 were obtained from the initial BRV outbreak. Samples from 2022 and 2024 were collected from animals found to be exhibiting clinical signs consistent with BRV during annual population monitoring surveys. Swabs from each turtle were collected from the buccal cavity, conjunctiva and cloaca and individually placed into 3mL of phosphate buffered gelatine saline (PBGS, pH 7.3) and held at 4°C prior to transport to the virology laboratory at the Elizabeth Macarthur Agriculture Institute for BRV and turtle serpentovirus RT-qPCR. Post-mortem samples were collected and frozen at -80°C immediately after euthanasia of affected turtles including brain or spinal cord, conjunctiva (eyelid), gonad, heart, kidney, liver, lung, rectum, and spleen (Supplementary Table 1). Additional representative portions of each organ were fixed in 10% neutral buffered formalin, processed in ethanol, embedded in paraffin, sectioned, stained with haematoxylin and eosin, mounted with a cover slip and examined by light microscopy.

### Sample extraction and total RNA-sequencing

Tissue samples were homogenised in RLT Plus buffer with beta-mercaptoethanol using TissueRuptor (QIAGEN, Hilden, Germany). RNA was then extracted using QIAGEN RNeasy Plus Mini Kit (QIAGEN, Hilden, Germany) following the manufacturer’s protocol. An additional negative control was established by passing extraction buffer through clean kit columns to monitor for contamination introduced during the extraction process. A total of 142 samples was extracted and pooled into 17 libraries, based on year of collection and tissue type, as well as the negative control. Pooled libraries were then transported to the Australian Genome Research Facility (AGRF, Melbourne, Australia) for total RNA sequencing. RNA pools were assessed for DNA contamination using a parallel capillary electrophoresis instrument (TapeStation; Agilent, California, USA), and quantified using the Qubit fluorometer (ThermoFisher Scientific, Massachusetts, USA). Pooled RNA libraries were subjected to ribosomal RNA depletion using the Illumina rRNA depletion kit, prepped with the Illumina Truseq Total RNA Library Preparation Protocol, and sequenced on the Illumina NovaSeqX platform (150bp paired end; Illumina, California, USA).

### Quality control and *de novo* assembly

Adapters and poor-quality raw reads were trimmed using Trimmomatic (v.0.35) (12) with the flags ‘LEADING:3 TRAILING:3 MINLEN:36’. Read quality and successful trimming was verified using FastQC (v.1.9) (13). To validate host identity and to check for potential library contamination, taxonomic assignment of trimmed paired-end reads was performed using Kraken2 (v.2.1.3) (14) with the confidence flag set at 0.2 and the core nucleotide database (core_nt; as of June 2025). Kraken2 output was viewed using the Pavian program (15).

Trimmed paired-end reads were then *de novo* assembled into contigs using MEGAHIT (v.1.2.9) (16).

### Virus discovery pipeline and abundance quantification

Assembled contigs were screened using DIAMOND BLASTx (v.2.16.0; flags set at “-e 1E-5 --ultra-sensitive”) (17) against the Reference Virus Database (RVDB; v31) (18) and RdRp-Scan (19) databases. False positive virus contigs were further identified by screening against the NCBI non-redundant (nr) protein database (as of December 2025) (20). Putative virus contigs were subject to open reading frame (ORF) prediction using Geneious Prime

(v.2025.1) and a manual web-based BLAST to confirm virus-like contigs. A total of 62 virus contigs (with a minimum length >699bp) were identified in this study (Supplementary Table 2). Virus abundance was estimated using RNA-Seq by Expectation-Maximisation (RSEM) (21), with trimmed paired-end reads mapped to MEGAHIT assembled contigs and abundance reported in transcripts per million (TPM).

### Phylogenetic analysis

To place the virus sequences obtained here in phylogenetic context, a total of 30 background viruses was selected from NCBI Taxonomy and GenBank to represent the diversity within the *Tobaniviridae*. Phylogenetic trees were then estimated using amino acid alignments of three proteins – the RNA-dependent RNA polymerase (RdRp), ORF1ab, and the spike protein – with all sequences aligned using the MAFFT-E-INS-i algorithm (22), and trimmed for ambiguously/poorly aligned regions using trimAl (23). Maximum likelihood (ML) phylogenetic trees for these sequence alignments were then estimated using IQ-TREE2 (24) with the ‘ModelFinder Plus’ (-m MFP) flag used to select the best-fit model (25) and with 1000 bootstrap replicates. The resulting trees were annotated in FigTree (v1.4.4) (26), and edited using the *ggtree* package (v3.8.2) in R (v4.3.3) and Adobe Illustrator (2025).

### Primer design, RT-PCR, and Sanger sequencing

Primers were designed to amplify a 430bp region encompassing the predicted ribosomal frameshift (slippery) site between ORF1a and ORF1b of BRV2 (forward primer 5’-TTTTTCGAGGCGTTTGATTC-3’; reverse primer 5’-CAACCGGCTTTACAACATAC-3’). Three libraries (2015 kidney, 2022 lung, 2024 lung) were subject to reverse transcription polymerase chain reaction (RT-PCR) using the BRV2 slippery site primers and Invitrogen SuperScript IV One-Step RT-PCR System (Thermo Fisher Scientific, Massachusetts, USA). An additional negative control containing the primers and RNase/DNase free water was established to check for primer binding and potential laboratory contamination. A total reaction volume of 25µL was loaded into SimpliAmp Thermal Cycler (Thermo Fisher Scientific, Massachusetts, USA) with a cycle set at 95°C for 3min, 40 cycles of 95°C for 60s, 55°C for 30s and 72°C for 30s, and a final extension at 72°C for 7 minutes. Easy Ladder I (Meridian Bioscience, Ohio, USA) and PCR product with gel loading solution (Sigma-Aldrich, Massachusetts, USA) was loaded into 0.1% Agarose gel + SBYR Safe DNA Gel Stain (Thermo Fisher Scientific, Massachusetts, USA). The molecular imaging GelDoc XR+System (Bio-Rad, California, USA) confirmed the expected 430bp band. The PCR product was Sanger sequenced at the Ramaciotti Centre for Genomics (UNSW, Sydney, Australia) to confirm the presence of the canonical 5’-AAAAAAC-3’ slippery sequence and the adjacent RNA secondary ‘kissing loop’ structure (2, 27).

## RESULTS

### Pathology Summary

Fifteen adult Bellinger River snapping turtles (11 male, 4 female) were examined from the 2015 mortality event, five subadult turtles (3 female, 2 undetermined sex) were examined from the 2022 event, and 21 turtles (2 juveniles, 17 subadults and 2 adults; 10 male, 9 female, 2 sex undetermined) were examined from the 2024 event. Signalment, clinical and pathological findings from each animal are described in Supplementary Table 3 and summarised in Supplementary Table S4. Clinical signs were consistent across each event.

Affected turtles had mild to marked chemosis and were lethargic or moribund. In most circumstances, eyelids were so swollen that the eyes could not be visualised. Eyelid epithelium was frequently eroded to ulcerated. A small subset of animals had serous to fibrinoid ocular or nasal discharge, focal plaques covering the epidermis of the ventral hind limbs and tail, corneal oedema, keratitis, were pawing at their eyes, or exhibited hind limb paresis. The most common cause of death was euthanasia, although nine animals from the 2024 event were either found dead or were treated and subsequently died, and a proportion of animals was treated and survived infection in 2024.

Consistent lesions observed during gross post-mortem examinations conducted in 2015 included severe chemosis, miliary tan splenic foci, generalised or radiating patterns of renal cortical pallor with additional punctate tan foci, tan fibrinoid plaques on the skin of the ventral hind limb and tail, and fewer animals with oral tan foci or petechiae, hydropericardium, serosal oedema or hydrocephalus (Figure 1). Observations were similar in 2022 and 2024 (Figure 2), although these animals also had a high prevalence of splenomegaly, pulmonary consolidation, hydropericardium, serosal oedema, tan foci on serosal surfaces (spleen, pancreas, epicardium, intestine, pleura) and a low prevalence of renal pallor and exudative bronchopneumonia (n=1).

Common histological findings included a range of focal and mild, to marked and extensive fibrino-necrotising and granulocytic inflammation in the eyelid, conjunctiva, lacrimal gland, periocular soft tissues, cornea, anterior chamber of the globe, renal cortical interstitium, nasal sinuses and the wall of the urinary bladder. Vasculitis and intravascular thrombi were often multifocally evident within these inflammatory lesions and in otherwise normal tissues. Across each event there were animals with focal inflammation in gonads, the wall of the urinary bladder, oral mucosa, oesophagus, gastrointestinal tract, olfactory and optic nerves, meninges and brain (particularly the olfactory lobe), and less commonly, in skeletal muscle, myocardium, oviduct, thymus, and coelomic serosal membranes. Effacement of the lacrimal gland architecture was consistent across events. In 2015 there was also effacement of splenic parenchyma and renal collecting duct epithelium. During 2022, and through most of the 2024 events, the splenic tissue was spared, and hepatic and renal lesions most often had an embolic pattern, although mild, acute necrosis of collecting duct epithelium was observed. A higher proportion of animals in latter events also had fibrino-necrotising and granulocytic inflammation in the lower respiratory tract (trachea, pulmonary interstitium and bronchial epithelium).

**Fig. 1.**
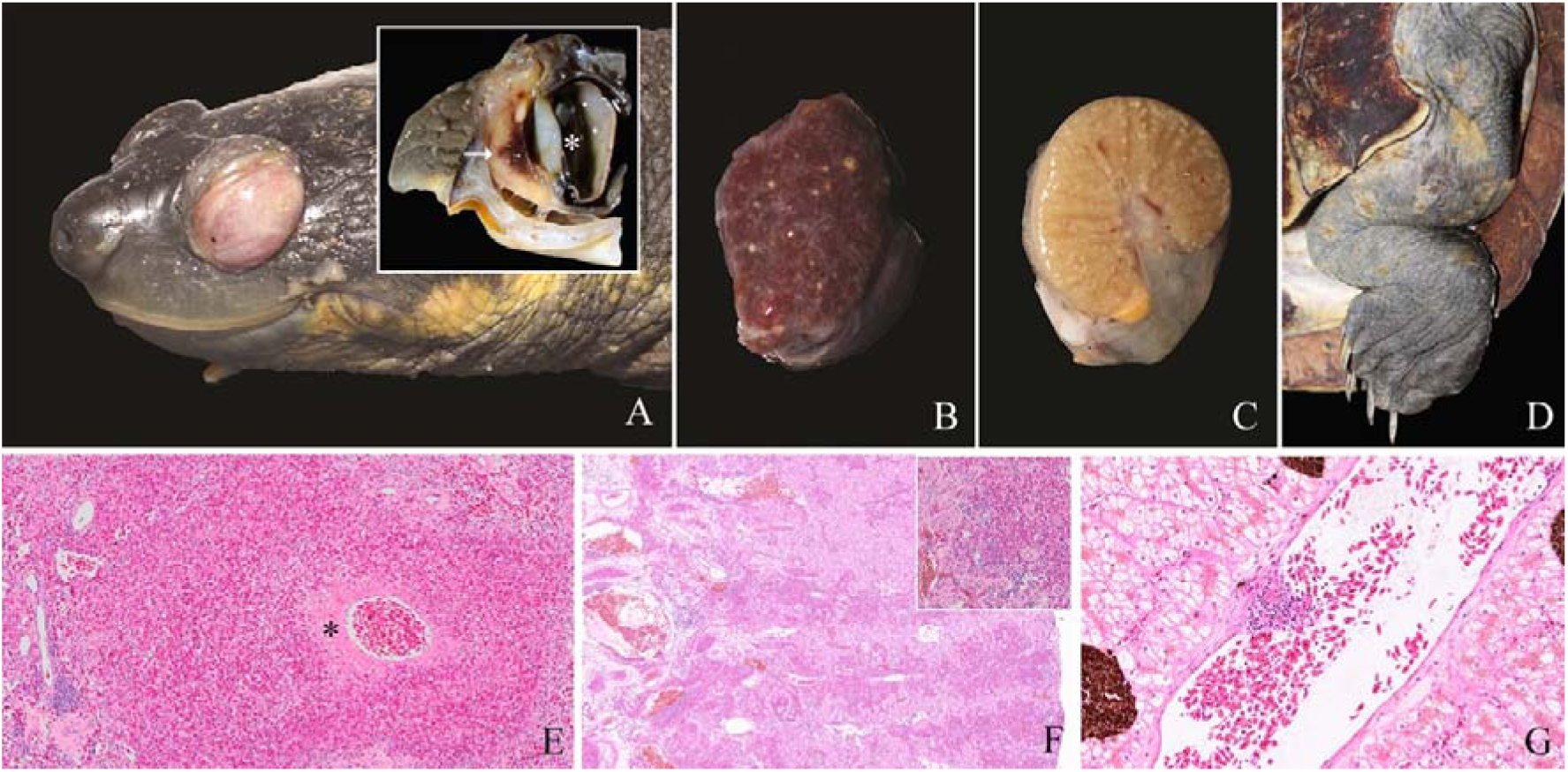
Gross and histological findings from the 2015 mass mortality of Bellinger River snapping turtles (*M. georgesi*). (A) Chemosis, with inset illustrating expansion and discoloration of the lacrimal gland (between arrows) and adjacent soft tissue and corneal oedema (asterisk). (B) Miliary tan splenic foci. (C) Generalised renal pallor, with alternating light and dark tan linear rays throughout the cortex. Miliary discrete, round, white to tan foci in the hilar connective tissue and cortex. (D) Multifocal epidermal erosions and ulcerations covered with tan exudate on the skin of the thigh and tail. (E) Kidney: Extensive splenic perivascular coagulation necrosis, fibrin deposition (asterisk) and granulocytic infiltrates. (F) Kidney: Linear rays of intact proximal renal tubules alternating with rays of intense interstitial granulocytic infiltration (inset) effacing collecting ducts. (G) Liver: Focal hepatic vasculitis characterised by mural and luminal fibrin and leucocytes.

Histological observations in a small subset of animals, which were interpreted as incidental to the investigation, included intestinal coccidiosis, renal tubular intracellular protozoa (interpreted as likely myxozoa), urinary bladder and renal tubular flagellate protozoa, urate nephrosis and protein losing nephropathy. From 2022 onwards, foreign body granulomas containing invertebrate matter morphologically consistent with processionary caterpillar (*Ochrogaster* sp.) setae were observed in oral and gastrointestinal mucosa, and less commonly in splenic capsule, mesentery and urinary bladder wall.

**Fig. 2.**
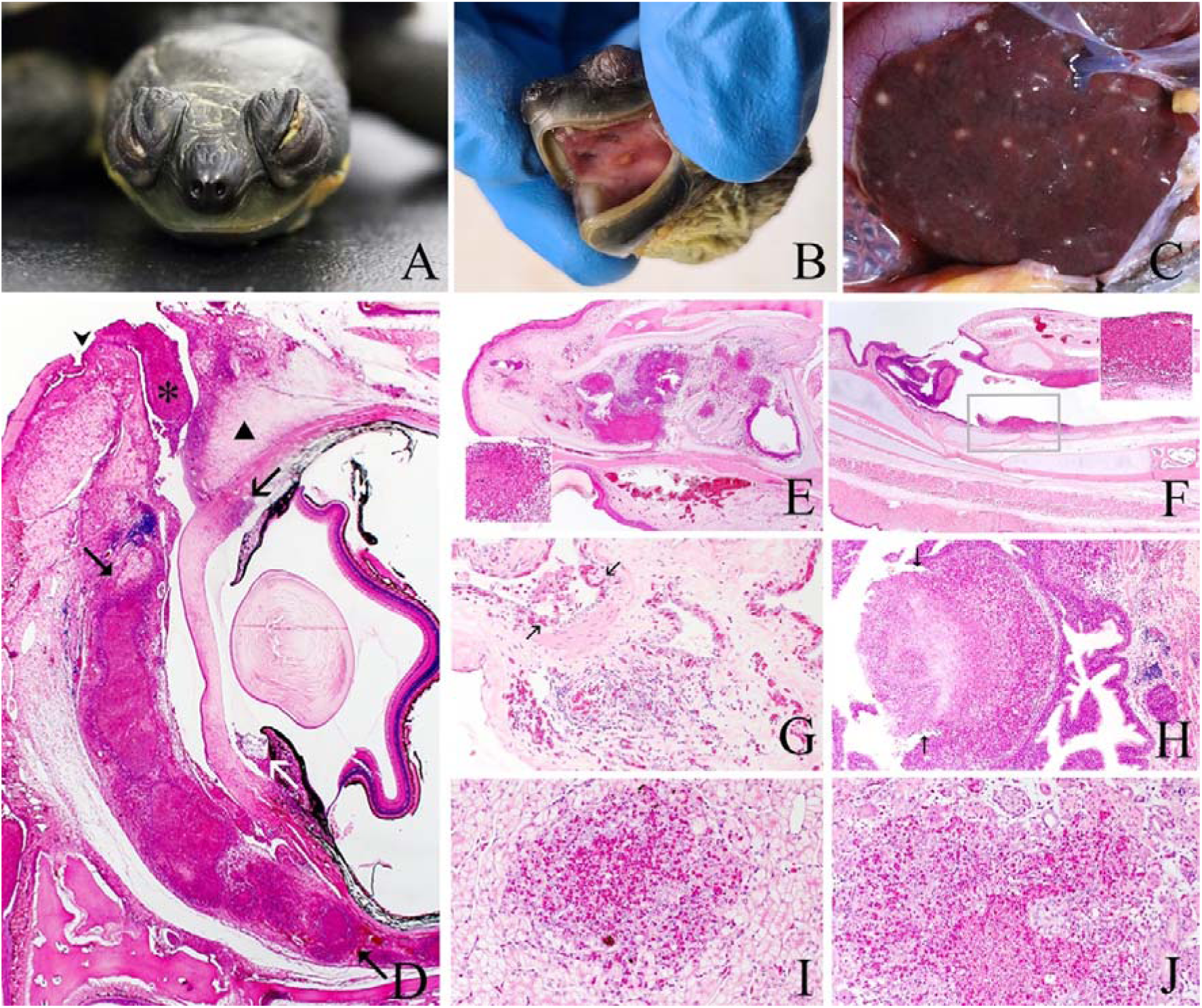
Gross and histological findings from the 2022 and 2024 mortality of Bellinger River snapping turtles. (A) Bilateral chemosis and unilateral tan ocular discharge. (B) Focal palatine lesion with tan centre and peripheral erythema. (C) Miliary tan hepatic foci. (D) Histological representation of ocular and periocular lesions common to all mortality events. Diffuse necrosis and granulocytic infiltrates in the lacrimal gland (between closed arrow). Oedema, fibrin deposition, and granulocytic infiltrates throughout the eyelids (triangle). Ulceration of the eyelid and conjunctival epithelium (arrowhead). Granulocytic exudate in the conjunctival sac (asterisk). Oedema and granulocytic infiltrates in the cornea and anterior ocular chamber (black and white open arrows respectively). (E) Sinuses: Multifocally extensive necrosis and severe granulocytic infiltration (inset) and exudation in the nares and sinuses. (F) Trachea: Multifocal tracheal mucosal necrosis and granulocytic infiltrates. (G) Lung: Necrosis of the bronchial epithelium (between arrows), interstitial oedema and granulocytic infiltrates. (H) Small intestine: segmental mucosal effacement, oedema and granulocytic infiltrates. (between arrows) (I) Liver: Focal hepatic necrosis with granulocytic infiltrates. (J) Kidney: focal necrosis with granulocytic infiltrates.

### Total RNA-Seq data summary

Total-RNA sequencing resulted in a total of 1,224,439,296 trimmed paired-end reads, equating to 427Gb of data across 16 sample libraries that range between 39,421,822-129,626,750 reads and average 76,527,456 reads per library. The negative control comprised 1,694,417 trimmed paired-end reads and 1.1Gb of data. Kraken2-based taxonomic classification of the sample libraries and negative controls reported no human contamination or unexpected bacterial or parasite signals within the sequencing data. No viruses were detected in the negative control library using the virus discovery pipeline, thereby excluding viral contaminants. As expected from tissue RNA extraction, *Testudines* (i.e., turtle) host reads were the most abundant order in every sequencing library, ranging between 81.1-99.8% and averaging 98.8% of reads at the order level.

### Detection of Bellinger River virus 1 (BRV1)

Herein, for clarity, the original BRV identified in 2015 will be referred to as Bellinger River virus 1 (BRV1) and the novel Bellinger River virus characterised in this study (see below) referred to as Bellinger River virus 2 (BRV2). Collectively, these viruses are referred to as the Bellinger River viruses (BRVs). BRV1 was detected in all 2015 *M. georgesi* libraries (kidney, liver, lung) with the highest abundance of 1061.33TPM in the 2015 kidney library (Figure 3). A complete 29,951bp BRV1 genome was extracted from the kidney library and compared against the reference BRV1 isolate J248 (GenBank accession MF685025.1). Phylogenetic analysis revealed that the virus detected in this study from archival 2015 samples and the independently identified BRV1 isolate J248 reference (2), share a 100% RdRp amino acid identity, 99.7% spike nucleotide identity and 98% full genome nucleotide identity (Figure 4). The detection of BRV1 in the 2015 samples also serves as a positive control for the metatranscriptomic pipeline, confirming its ability to detect and assemble BRV genomes. Strikingly, however, we did not detect BRV1 in any of the 2022 or 2024 samples.

**Fig. 3.**
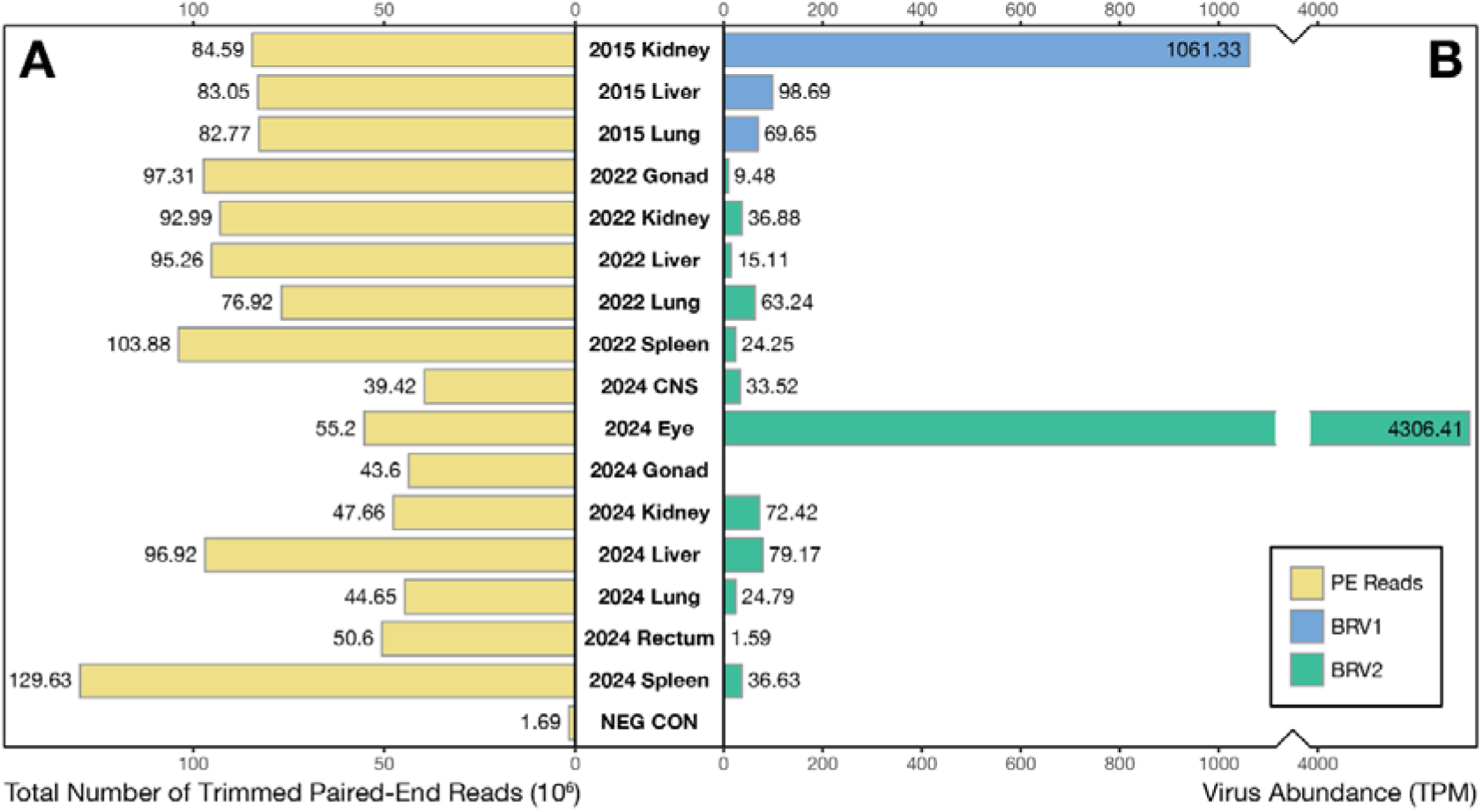
Summary of Bellinger River snapping turtle sequencing libraries. (A) Total number of trimmed paired-end reads in millions per sequencing library. (B) Virus read abundance measured by total the number of transcripts per million (TPM) in each library. Adjacent key represents the following: Yellow, paired-end reads; Blue, Bellinger River virus 1 (BRV1); Green, Bellinger River Virus 2 (BRV2).

### Detection of Bellinger River virus 2 (BRV2)

BRV2 was detected in all libraries from 2022 (gonad, kidney, liver, lung, spleen, central nervous system, conjunctiva) and in most libraries from 2024 (kidney, liver, lung, rectum and spleen). Notably, BRV2 was not detected in any 2015 samples. In the 2024 samples, the highest observed virus abundance was observed in conjunctiva (4306.41 TPM), liver (79.17 TPM) and kidney (72.42 TPM) libraries, while in 2022 the lung (63.24 TPM), kidney (36.88 TPM) and spleen (24.25 TPM) samples had the highest observed virus abundances (Figure 3). No BRVs were detected in 2024 gonad samples, although BRV2 was detected in the 2022 gonads (69.48 TPM). Full genomes of BRV2 were identified in 2022 lung (30,338bp) and 2024 lung (30,257bp). The BRV2 sequences from 2022 exhibited 99.8% RdRp amino acid identity, 98% spike nucleotide identity, and 98% full genome nucleotide identity with the BRV2 sequences detected in 2024. Together, these sequences represent a distinct BRV lineage that shares 91.0% RdRp amino acid identity, 75.4% spike nucleotide identity, and 70.6% full genome nucleotide identity with BRV1 (Figure 4C). Comparative genomic analysis of BRV1 and BRV2 revealed similar genomic organisation (Figure 4A) with no evidence of genome recombination with BRV1. Mutations that differentiated the two viruses were present across the viral genome, further supporting the interpretation that BRV2 represents a genetically distinct lineage. As expected, phylogenetic analysis placed BRV1 and BRV2 as sister lineages, which we can preliminarily classify as distinct virus species within the *Serpentovirinae* (Figure 4B, Supplementary Figure S1).

**Fig. 4.**
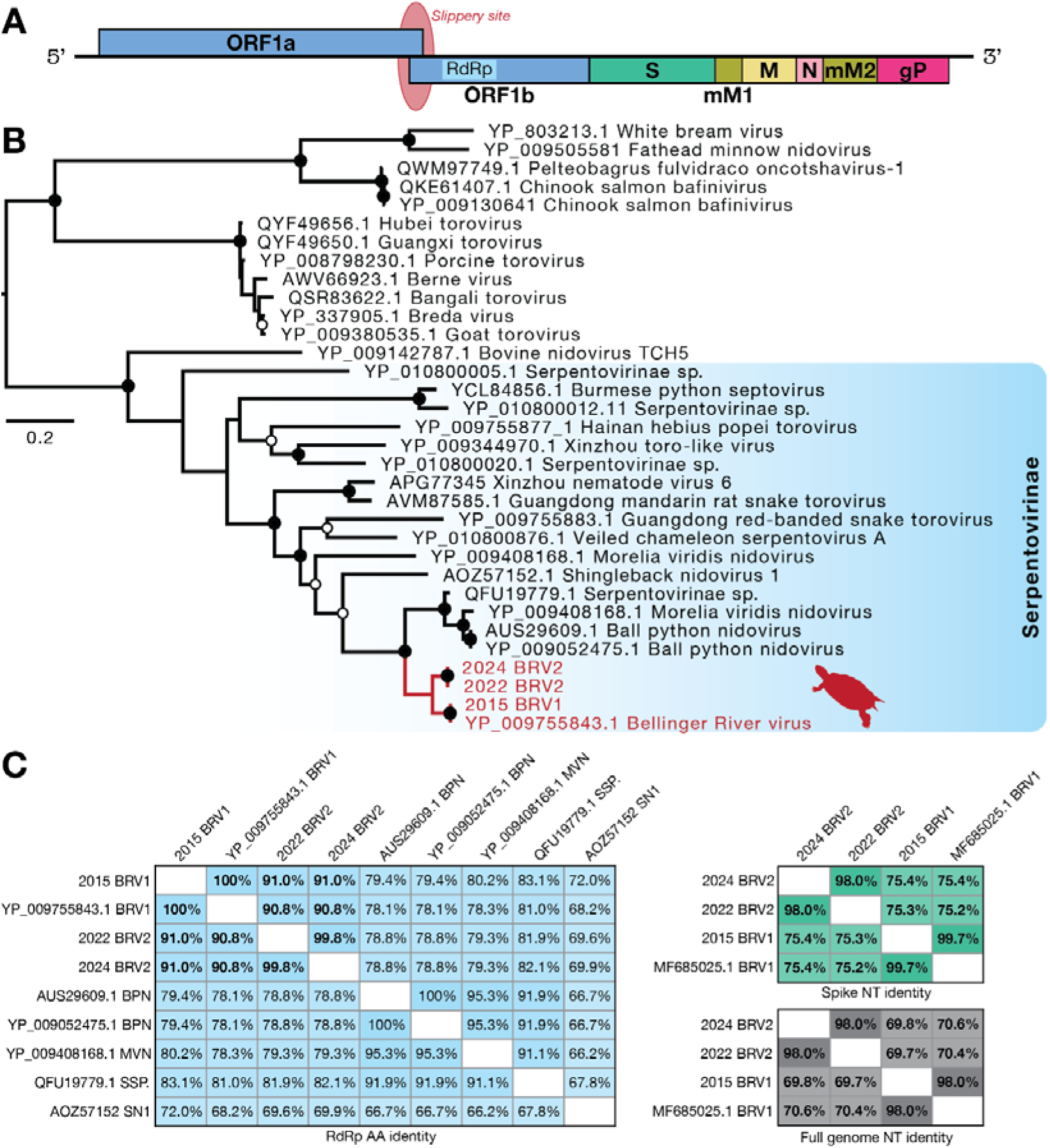
Evolutionary relationships of Bellinger River viruses, BRV1 and BRV2. (A) Schematic representation of the Bellinger River virus genome. (B) Maximum likelihood phylogenetic tree of the *Tobaniviridae* based on RdRp amino acid sequences. BRV sequences are marked in red. Node support was estimated using 1000 bootstrap replicates. The tree is mid-point rooted with nodes ≥95% bootstrap support annotated with black circles (●) and those 70-94% bootstrap support with white circles (○). The scale bar represents number of estimated amino acid substitutions per site. (C) Similarity matrices showing BRV RdRp amino acid identity in light blue, spike nucleotide identity in green, and full genome nucleotide identity in grey. Abbreviated virus names: Bellinger River virus 1 (BRV1), Bellinger River virus 2 (BRV2), Ball python nidovirus (BPN), Morelia viridis nidovirus (MVN), Serpentovirinae sp. (SSP.), Shingleback nidovirus 1 (SN1).

### PCR and the confirmation of slippage site

PCR of the 2024 lung, 2022 lung, and 2015 kidney libraries was conducted to confirm the slippage site between ORF1a and ORF1b. This slippage site, confirmed as a 5’-AAAAAAC-3’ sequence, allows for slippery binding of ribosome to ORF1ab template so that ORF1b, which contains the RdRp, can be independently translated. As expected, the 2015 sample was PCR negative for the 430bp band using the BRV2-specific primers, while all the 2022 lung and 2024 lung libraries were positive for BRV2. Negative controls only comprising primer water similarly revealed no bands. Sanger sequencing of the PCR amplicon further confirmed the slippage site and “kissing loop structure” by comparing homologous sequences (2). This site of ribosomal frameshift shared 100% sequence identity with the *de novo* assembled contig.

### Other viruses detected in Bellinger River snapping turtles

As well as the BRVs, we used metatranscriptomic sequencing to determine whether these turtle samples harboured additional viruses. Accordingly, a bastrovirus and a member of the *Cystoviridae* were detected in the 2024 *M. georgesi* samples with abundances of 0.55 TPM in spleen and 0.35 TPM in rectal tissue, respectively. No other viruses were detected. As the *Cystoviridae* are known to infect gram-positive bacteria rather than vertebrates (28), the putative cystovirus contigs detected in the rectal library were assumed to represent bacteriophage and hence excluded from further analysis. Bastroviruses are an unclassified group of non-enveloped, single-stranded, positive-sense RNA viruses that have been detected in human and animal faeces (29). Their genomes encode hepevirus-like non-structural proteins and astrovirus-like structural proteins. A 2351bp contig representing the RdRp of a novel bastrovirus was detected in spleen tissue and shared 94.4% amino acid identity with bastrovirus sp. (ASM79505.1), originally detected in Brazilian sewage (Supplementary Figure S2). Phylogenetic analysis placed this novel *M. georgesi* bastrovirus within a clade predominantly comprising environment-associated viruses with no evidence of vertebrate infection. Additionally, *M. georgesi* bastrovirus was only found in one library and at relatively low abundance to BRV2 (0.55TPM bastrovirus; 36.33TPM BRV2). Hence, the detection of this bastrovirus is likely incidental and requires further investigation beyond the scope of this study to determine the host range of these divergent bastroviruses.

## DISCUSSION

The Bellinger River snapping turtle has been the focus of more than a decade of conservation efforts. Following the mass mortality event in 2015, ongoing surveillance of *M. georgesi* health, population monitoring, and captive breeding efforts have supported the persistence of the extant population (1). Epizootics of Bellinger River snapping turtle morbidity and mortality in 2022 and 2024 were accompanied by the hallmark ocular and lacrimal gland lesions associated with BRV1, but internal lesion distribution and intensity varied and BRV1 PCR tests returned negative results, inciting further investigation. Herein, we report the identification of a second novel nidovirus (*Serpentovirinae*) in Bellinger River snapping turtles, further emphasising the need for ongoing *M. georgesi* health surveillance, disease investigation and conservation management.

Our phylogenetic analysis suggests that although BRV1 and BRV2 clearly share a common ancestor, forming a distinct BRV clade within the *Serpentovirinae*, they are sufficiently divergent in genome sequence (∼30% at the full genome nucleotide level) that BRV2 is unlikely to have directly emerged from BRV1 during the course of these outbreak investigations. Although there is insufficient data to perform molecular clock dating to obtain the timescale of virus evolution, the observed level of sequence divergence between BRV1 and BRV2 is too substantial to have been generated solely by *in situ* evolution during the seven-year period between 2015 and 2022. As such, we suggest that BRV2 emerged independently to infect *M. georgesi*. From the data collected in this study, we report that BRV1 was only detected in 2015 samples and absent in 2022 and 2024 samples, such that it is possible BRV2 has replaced BRV1 as the primary turtle pathogen in the Bellinger River.

Indeed, BRV1 has not been detected in routine surveillance of the Bellinger River since November of 2018 (5). Conversely, BRV2 could only be detected in the 2022 and 2024 samples and was absent in 2015 samples. However, these observations should be interpreted with caution as the limited number of samples in this study (n=35) may not accurately represent the prevalence and circulation of BRVs within the wild population.

Several non-mutually exclusive theories may explain these observations. First, the 90% population decline in 2015 likely caused a severe population bottleneck that reduced the number of susceptible adult hosts below the threshold required for sustained BRV1 transmission, which would be particularly apparent in an animal population that only spans a 60 km river. These epidemiological and ecological factors may account for the decline in BRV1 prevalence in samples collected after the 2015 outbreak. Other possibilities include changes in host immunity or the persistence of BRV1 at prevalences below the detection limits in this study. While our data suggests that BRV2 emerged independently to infect *M. georgesi*, it is unclear whether the appearance of this virus occurred after the decline of BRV1 or co-circulated with BRV1 while evading BRV1-specific molecular detection assays, before ultimately becoming the dominant circulating virus. In either scenario, the sudden emergence of BRV2 in *M. georgesi* is compatible with a recent cross-species transmission event from an unidentified reservoir host within the Bellinger River ecosystem. Determining the reservoir host of both BRVs will require broader surveillance of freshwater turtles, reptiles and other aquatic vertebrates within the Bellinger River catchment. This surveillance should include exotic or introduced hosts such as abandoned captive and commercially bred turtles and snakes, and reptiles translocated through rehabilitation or community action. As *E*. *macquarii* has also tested positive for BRV1 (5), the prevalence of BRV2 in *E*. *macquarii* should similarly be tested in future disease surveillance. Continued longitudinal surveillance using diagnostics capable of detecting both BRV1 and BRV2 will be necessary to determine whether BRV1 has truly been lost in the wild population or continues to circulate at low prevalence in *M. georgesi* or another host in the Bellinger River ecosystem. Turtle serpentovirus RT-qPCR testing should also be considered to assist in the identification of future viral spill-over events.

Also of note was that BRV2 was most abundant in conjunctiva tissue samples – some three orders of magnitude greater than in any other BRV2-positive library. This is consistent with previous observations that conjunctival swabs offered the most optimal sampling type for ongoing BRV1 surveillance (5). Conjunctival swabs may therefore provide a minimally invasive way to sample *M. georgesi* for BRVs. Additionally, rectal tissue provided a relatively low abundance of BRV2 which may indicate limitations in cloacal swab sampling for BRV surveillance, although further study is needed. The primers developed in this study, or newly designed primers targeting the BRV2 spike protein based on the genome sequences obtained here, may provide a cost-effective approach for the targeted molecular surveillance of BRV2, reducing the need for further metatranscriptomic sequencing. Incorporating turtle serpentovirus RT-qPCR and BRV2 specific PCRs into ongoing health monitoring programs will also improve the early detection of future BRV2 outbreaks and enhance the understanding of its prevalence, distribution, and epidemiology within the wild population.

This approach enables the monitoring of BRV2 circulation within the *M. georgesi* population and the potential future emergence of another novel serpentovirus.

In sum, this study characterises BRV2 as a novel virus that is potentially associated with negative health outcomes in *M. georgesi*, and illustrates the value of thorough and unbiased approaches to wildlife disease surveillance. The integration of total RNA sequencing with pathology and existing molecular diagnostic assays for BRV1 enabled the identification and characterisation of BRV2 and the exclusion of other potential co-pathogens. Continued pathogen surveillance will be critical to inform disease management and conservation in *M. georgesi*. However, experimental infection studies of BRV2 in *M. georgesi* are required to confirm virus replication and pathogenicity, although our study supports an aetiological link as BRV2 is the only candidate pathogen detected in the sequencing data. As low genetic diversity exists in its extant and endangered wild population (30), *M. georgesi* remains especially vulnerable to emerging infectious diseases.

## Acknowledgements

The Australian Registry of Wildlife Health (the Registry) is a 40-year science program of Taronga Conservation Society, which contributes to the surveillance of marine species health in partnership with the New South Wales Departments of Primary Industries and Regional Development and Climate Change, Energy, the Environment and Water. We thank the previous and ongoing efforts of staff and volunteers in assisting with managing samples and data related to wildlife health programs, particularly Drs. Peter Johnson and Bryn Lynar, and Nicole Dobson, Allison Rubie, and Karina Hammond. We also recognize the technical assistance provided by the Sydney Informatics Hub, a Core Research Facility of the University of Sydney.

## Author contributions

L.L., K.R. and E.C.H. conceived the study; K.R. and J.H. performed the animal sampling, histopathology and veterinary work; L.L., B.J.L., and J.R. performed the experimental and computational analyses; L.L., B.J.L., J.R., K.P., J.H., K.R., and E.C.H wrote the manuscript.

