## Supplementary figures and images for "Bellinger River Virus 2: The Emergence of a Second Novel Pathogenic Nidovirus in Bellinger River Snapping Turtles (*Myuchelys georgesi*)"

### Supplementary Figures S1-S2

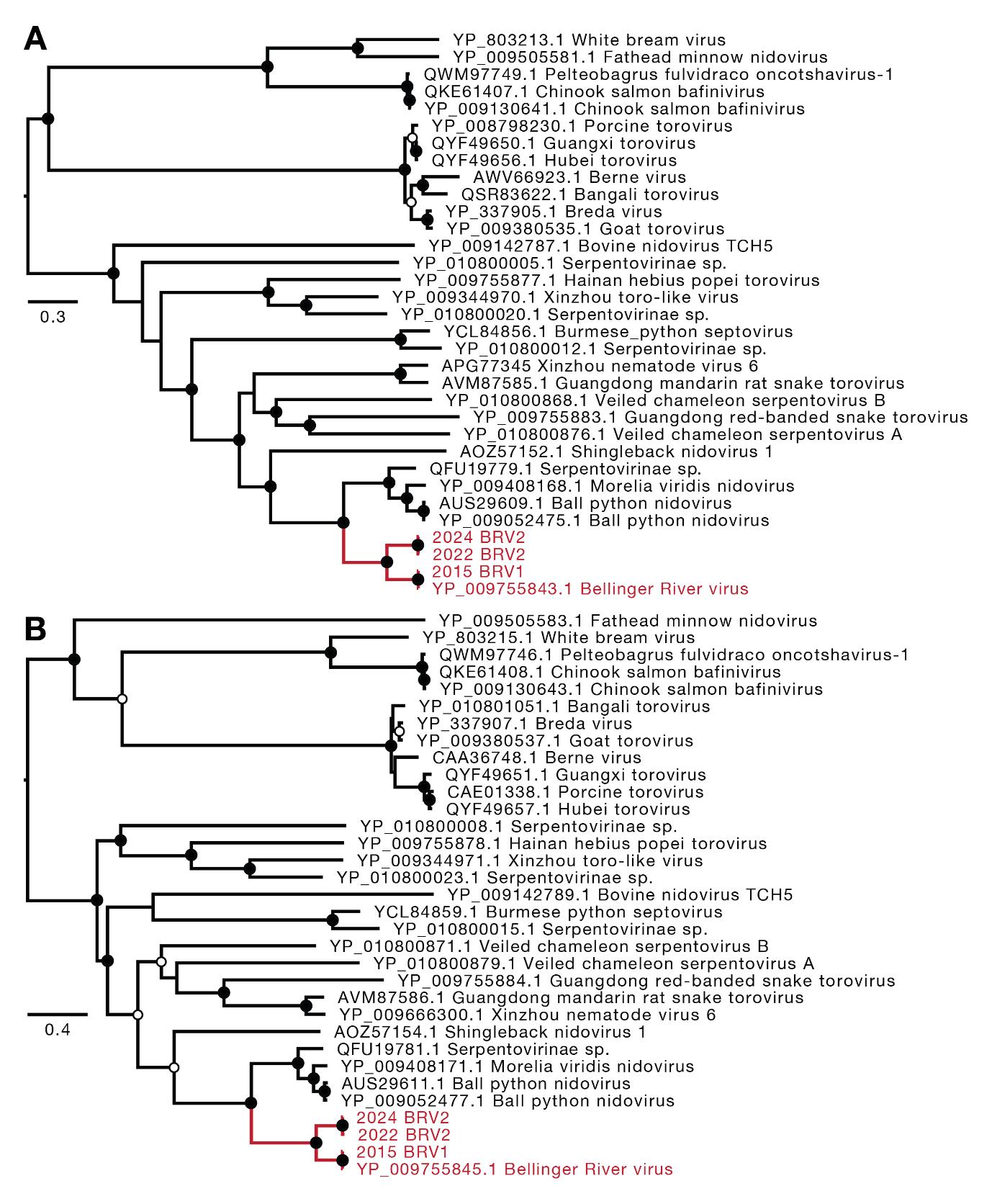

**Supplementary Figure 1**

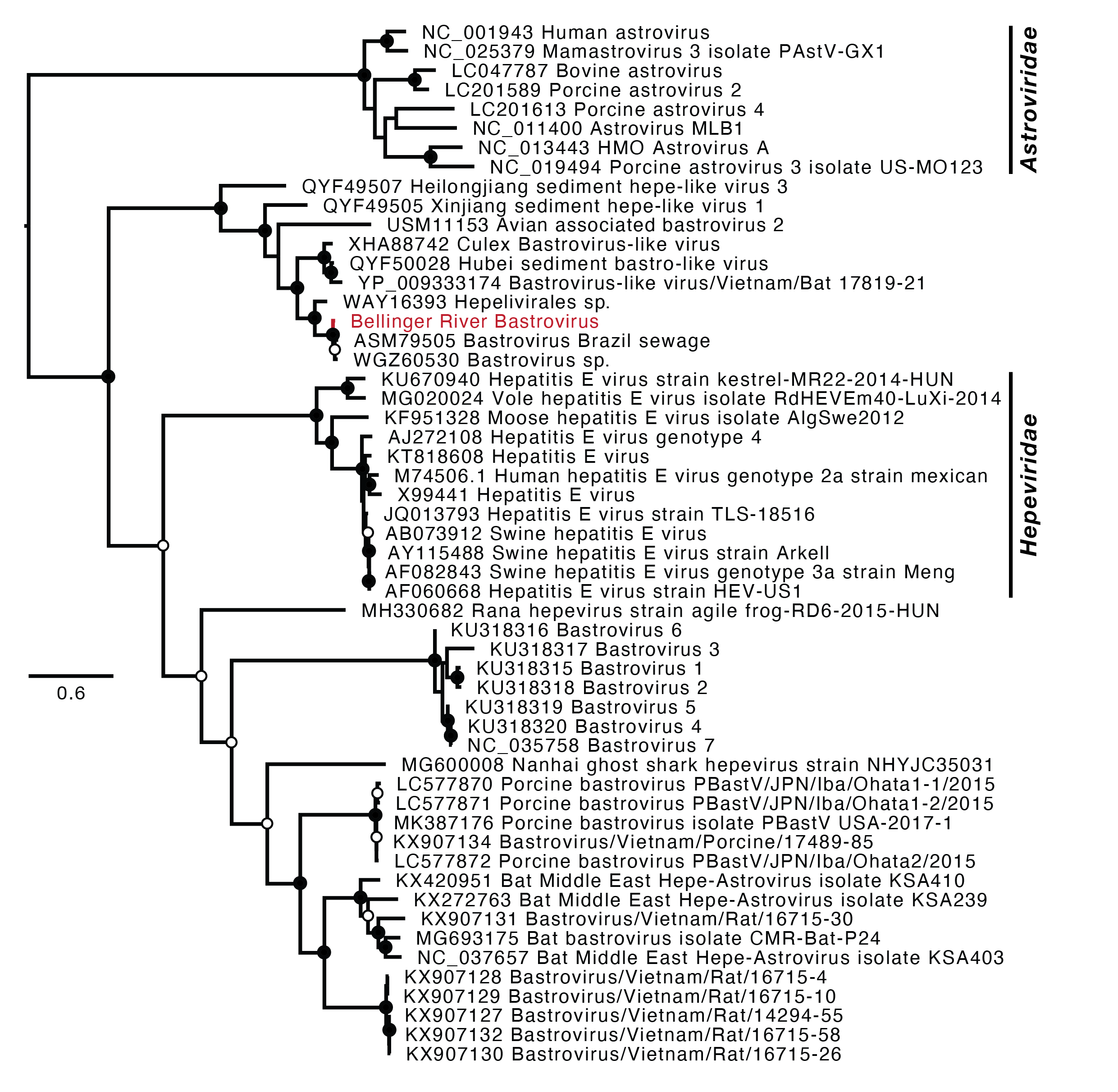

**Supplementary Figure 2**
