## Supplementary Tables S3 for "Bellinger River Virus 2: The Emergence of a Second Novel Pathogenic Nidovirus in Bellinger River Snapping Turtles (*Myuchelys georgesi*)"

| **Table S3**. Signalment and pathology findings in Bellinger River snapping turtles | | | | | | | | | | | | | | | | | |
| --- | --- | --- | --- | --- | --- | --- | --- | --- | --- | --- | --- | --- | --- | --- | --- | --- | --- |
| **Case**  **Number** | **CCL (mm)** | **Weight**  **(g)** | **Sex** | **Age class** | **Date** | **Euthanased/ Died** | **Gross lesions** | **Eyes** | **Spleen** | **Kidney** | **Bladder** | **Vasculitis** | **CNS/PNS** | **Respiratory** | **Liver** | **Digestive tract** | **Other** |
| 10446.1 | 175 | 469 | Male | Adult | 20-2-2015 | Euthanased | Bilateral eyelid swelling  Unilateral eyelid necrosis | Blepharitis and conjunctivitis – ulcerative, necrotising, granulocytic, LP, edema, fibrin, E, M3  Keratitis – granulocytic, ulcerative, E, M2  Anterior uveitis – granulocytic, multinucleate giant cells, D,M3  Dacryoadenitis – granulocytic, necrotising, dysplastic epithelium, D, M2  Periocular cellulitis - LP granulocytic, edema, osteolysis, E, M2  Choroiditis – granulomatous, MF, M1 | Splenitis - granulocytic, necrotising, fibrin, MFE, M2-3 | NA | Cystitis – granulocytic, edema, E, M1 | Intravascular thrombi: spleen, conjunctiva, | Olfactory and optic neuritis -macrophages and granulocytes, E, M2  Cerebral lipofuscinosis – MF, M1 | Pleural edema – D, M1 | Focal portal edema | Colitis with mural edema – fibrino-necrotising, granulocytic, granulomatous, E, M2  Gastric mural thrombosis, granulocytic, LP, edema, F, M1 | Testicular spermatogenesis  Epidermitis and dermatitis – granulocytic, edema, F, M1 (hind leg)  Pancreatitis – LP, MF, M1 |
| 10446.2 | 157 | 332 | Male | Adult | 20-2-2015 | Euthanased | Bilateral eyelid swelling  Unable to see corneas | Blepharitis and conjunctivitis – ulcerative, necrotising, granulocytic, edema, fibrin, E, M3  Keratitis – granulocytic, ulcerative, E, M2  Anterior uveitis – granulocytic, necrotising, D,M3  Dacryoadenitis – granulocytic, necrotising, dysplastic epithelium, D, M2  Periocular cellulitis - granulocytic, LP, edema, E, M2 | Splenitis – granulocytic - granulocytic, necrotising, fibrin, thrombi, E, M2-3 | NA | Cystitis – granulocytic, necrotising, fibrin, thrombi, E, M1 | Intravascular thrombi: spleen, kidney, lungs, small intestine, oral | Meningoencephalitis – E, M2  Olfactory and optic neuritis – MRE, M2  Cerebral lipofuscinosis, MF, M1 | Broncho-interstitial pneumonia – granulocytic, thrombi – MF, M1-2  Pleural edema – E, M1 | Hepatitis – granulocytic, MF, M1 | Oral mucosal edema and thrombi – MF, M2  with lamina proprial thrombi  Gastroenteritis – E, M1 | Testicular spermatogenesis  Hydrocephalus  Myodegeneration, myositis – granulocytic edema or fibrin, E, M2 |
| 10446.3 | 178 | 501 | Male | Adult | 20-2-2015 | Euthanased | Bilateral eyelid swelling, erythema, clear discharge  Raised tan lingual lesion 1mm diameter | Blepharitis and conjunctivitis – ulcerative, necrotising, granulocytic, edema, fibrin, E, M3  Keratitis – granulocytic, ulcerative, E, M2  Anterior uveitis – granulocytic, necrotising, D,M3  Dacryoadenitis – granulocytic, necrotising, dysplastic epithelium, D, M2  Periocular cellulitis - granulocytic, LP, edema, E, M2 | Splenitis – granulocytic, necrotising, fibrin, D, M2-3 | Interstitial nephritis –granulocytic, necrotising, fibrin, thrombi, fibrin, MFE, M3 | Cystitis – granulocytic, LP, edema, D, M1 | Intravascular thrombi: liver, spleen, testis, bladder, kidney, intestinal serosal, oral mucosa, epididymis | Meningitis – granulocytic, macrophages, hemorrhage, fibrin, E, M1  Cerebral lipofuscinosis – MF, M1 | Interstitial vasculitis with mural edema and granulocytes extending into adjacent parenchyma. Adjacent resp epithelium desquamated with lost cilia – MF, M1 | Portal vein thrombosis -MF, M1  Cholecystitis – granulocytic, edema, F, M1 | Oral ulceration - F, M1  Gastric edema – macrophages, E, M1 | Testicular spermatogenesis,  Interstitial orchitis - granulocytic, thrombi, MF, M2 |
| 10446.4 | 219 | 930 | Female | Adult | 20-2-2015 | Euthanased | Bilateral eyelid swelling, erythema, lower eyelid ulceration with fibrinoid exudate.  3-4 Raised tan lingual lesions 1mm  Liver large and friable.  Miliary white foci splenic parenchyma  Bladder wall thick | Blepharitis and conjunctivitis –granulocytic, necrotising, edema, fibrin, E, M3  Keratitis – granulocytic, edema, E, M2  Anterior uveitis – granulocytic, D, M2  Dacryoadenitis – granulocytic, necrotising, D, M2  Periocular cellulitis - granulocytic, LP, edema, E, M2 | Splenitis - granulocytic, necrotising, fibrin, thrombi, MF, M2, extending into capsule | Interstitial nephritis, granulocytic, fibrin, thrombi, MF, M2-3  Renal tubular and collecting duct necrosis - MF, M1-2 | Cystitis – ulcerative, granulocytic, LP, edema, necrosis, D, M2 | Intravascular thrombi: meninges, renal, oviduct, stomach, spleen | Meningitis – granulocytic, thrombus, MF M1  Cerebral lipofuscinosis - MF, M2 | Sinusitis – granulocytic,  fibrinonecrotising, edema, MF, M2  Interstitial pneumonia – granulocytic, edema, vasculitis with mural edema MF, M2  Respiratory epithelial desquamation and necrosis, MF, M1  Pleural edema, D, M1 | Hepatic subcapsular hemorrhage – granulocytic, necrosis, F, M2 | Stomatitis- fibrinonecrotising, MF, M2  Gastric wall thrombi MF, M1  Small and large intestine – granulocytes, LP edema, D, M1 | Oviduct – granulocytic inflammation lamina proprial, serosal thrombi, MF, M2  Interstitial oophoritis – granulocytic, necrotising, fibrin – MF, M2 - extending into coelomic cavity  Epicarditis – granulocytic, fibrin, MF, M1 |
| 10446.5 | 169 | 398 | Male | Adult | 20-2-2015 | Euthanased | Bilateral lower eyelid swelling and epidermal ulceration | Blepharitis and conjunctivitis – ulcerative, necrotising, granulocytic, edema, fibrin, E, M3  Keratitis – ulcerative, granulocytic, necrotising, edema, E, M3  Anterior uveitis – granulocytic, macrophages, D, M2  Dacryoadenitis – granulocytic, necrotising, D, M2  Periocular cellulitis - granulocytic, LP, edema, E, M2 | Splenitis - granulocytic, fibrinonecrotising, thrombi, MFE-D, M2-3 extending into capsule | Interstitial nephritis – granulocytic, fibrinonecrotising, MF, M2  Collecting duct necrosis – MF, M1  Membranous glomerulopathy – MF, M1 | Cystitis – granulocytic, edema, thrombi, MF, M1-2 | Intravascular thrombi: pancreas, liver, bladder | NVL | Sinusitis – ulcerative, fibrinonecrotising and granulocytic, hemorrhagic, E, M3  Lung - NA | Hepatic intravascular thrombi – MF, M1 | Stomatitis - granulocytic, necrotising, MF, M2  Gastritis – granulocytic, serosa thrombi – MF, M1  Large intestinal ulceration – granulocytic, F, M1 | Testicular spermatogenesis  Orchitis – granulocytic, necrotising, F, M1  Coelomitis - pancreatic capsule – granulocytic, necrotising, F, M1  Thyroiditis – interstitial, granulocytic, necrotising, MF. M1  Epicardial edema , LP, MF, M1  Epidermal degeneration – F, M1  Dermatitis – granulocytic, edema, F, M1  Myodegeneration – F, M1 |
| 10446.6 | 173 | 483 | Male | Adult | 20-2-2015 | Euthanased | Bilateral lower eyelid swelling – focal unilateral tan 1 mm focus medial canthus of eye  Mildly pale renal parenchyma | Blepharitis and conjunctivitis – ulcerative, necrotising, granulocytic, edema, fibrin, E, M3  Keratitis – ulcerative, granulocytic, necrotising, edema, E, M3  Anterior uveitis – granulocytic, macrophages, FE, M1  Dacryoadenitis – granulocytic, necrotising, D, M2  Periocular cellulitis - granulocytic, LP, edema, E, M2 | Splenitis - fibrino-necrotising and granulocytic, MFE, M3 | Interstitial nephritis – linear cortical tracts of granulocytic, fibrino-necrotising, thrombi, MFE, M3 | Cystitis – granulocytic, LP, edema, D, M1 | Intravascular thrombi: spleen, kidney. | NVL | Sinusitis – granulocytic, fibrinonecrotising, E, M3 | NVL | Stomatitis – necrosis and edema, MF, M1  Gastritis, granulocytic, edema, MF, M1  Large intestinal edema, granulocytes, LP, MF, M1 | Spermatozoa in vas, no spermatogenesis  Interstitial orchitis – granulocytic, edema, F, M1  Adrenitis granulocytic, fibrino-necrotising - MFE, M3  Dermatitis – granulomatous, necrotising, F, M1 |
| 10446.7 | 176 | 543 | Male | Adult | 20-2-2015 | Euthanased | Bilateral eyelid swelling and erythema.  Lingual petechiae  Diffusely pale renal parenchyma | Blepharitis and conjunctivitis – ulcerative, necrotising, granulocytic, hemorrhage, edema, fibrin, E, M3  Keratitis – ulcerative, granulocytic, macrophages, necrotising, edema, E, M3  Anterior uveitis – granulocytic, macrophages, FE, M1  Dacryoadenitis – granulocytic, necrotising, D, M2  Periocular cellulitis - granulocytic, LP, edema, E, M2  Choroiditis – LP, fibrin, F, M1 | Splenitis - fibrino-necrotising and granulocytic – MFE, M3 | Interstitial nephritis – linear cortical tracts of granulocytic, firbrino-necrotising, thrombi, MFE, M3 | Cystitis – granulocytic, LP, edema, D, M1 | Intravascular thrombi – pleura, liver, | Optic neuritis – granulocytic, macrophages, F, M1  Cerebral lipofuscinosis – MF, M1 | Sinusitis - fibrinonecrotising and granulocytic, FE, M2  Pleural edema, thrombi, M1 | Hepatic necrosis – granulocytic, necrotising, fibrin, thrombi, MF, M1 | Oral mucosal necrosis – MF, M1  Large intestinal erosion, granulocytic, edema, MF, M1 | Testicular spermatogenesis  Myocardial necrosis and edema, intranuclear basophilic inclusion bodies, F, M1  Epidermal ulcer and dermatitis – granulocytic, F, M1 |
| 10446.8 | 168 | 398 | Male | Adult | 20-2-2015 | Euthanased | Bilateral eyelid swelling, hyphema left eye and tan discharge. Right lid erythematous and ulcerated.  Focal haemorrhage along choanal slit  Pale renal parenchyma | Blepharitis and conjunctivitis – ulcerative, necrotising, granulocytic, edema, fibrin, E, M3  Keratitis – ulcerative, granulocytic, macrophages, necrotising, edema, E, M3  Anterior uveitis – granulocytic, macrophages, FE, M1  Dacryoadenitis – LP, necrotising, D, M2  Periocular cellulitis - granulocytic, LP, edema, E, M2 | Splenitis – granulocytic and fibrinonecrotising, MFE, M3 | Interstitial nephritis – linear cortical tracts of granulocytic, firbrino-necrotising, thrombi, MFE, M3 | Cystitis – ulcerative, granulocytic, edema, MF, M1 | Intravascular thrombi, stomach, liver, jejunum, bladder | Hydrocephalus and cerebral oedema with ependymal fibrinoid material  Cerebral lipofuscinosis – MF, M1 | NVL | Sinusitis – LP, MF, M1 | Stomatitis – necrotising and granulocytic – F, M1  Gastric necrosis – fibrin, thrombi, MF, M1  Colitis – granulocytic, LP, necrotising, edema, D, M1 | Testicular spermatogenesis  Epidermal necrosis – focal, mild, acute  Pancreatic and hepatic atrophy – D, M1  Protozoa in urinary bladder and renal tubular lumens |
| 10446.9 | 179 | 500 | Male | Adult | 20-2-2015 | Euthanased | Bilateral eyelid swelling, lower more severe and ulcerated with tan fibrinoid exudate  Inguinal erythema  Miliary white splenic foci | Blepharitis and conjunctivitis – ulcerative, necrotising, granulocytic, edema, fibrin, E, M3  Keratitis – granulocytic, edema, E, M3  Anterior uveitis – granulocytic, fibrin, D, M2  Dacryoadenitis – granulocytic, necrotising, D, M2  Periocular cellulitis - granulocytic, LP, edema, E, M2 | NA | Interstitial nephrosis, - granulocytic, necrotising - MFE, M3 | Cystitis – granulocytic, necrotising, fibrin, MF, M1-2 | NVL | Cerebral lipofuscinosis – MF, M1 | Sinusitis – ulcerative, granulocytic, LP, necrotising, MF, M1-2  Lung - NA | Hepatitis- granulocytic, fibrinonecrotising – F, M1 | Stomatitis – granulocytic, necrotising, MF, M1-2  Enteritis – perivascular – granulocytic, necrotising, fibrin, MF, M1  Large intestinal mural, edema - D, M1  Gastric granuloma – F, M1 | Testicular spermatogenesis  Epidermal necrosis and intercellular edema– F, M1 |
| 10446.10 | 183 | 528 | Male | Adult | 20-2-2015 | Euthanased  Frothing at the mouth and gasping | Oral froth  Bilateral eyelid swelling - M1 | Blepharitis and conjunctivitis – granulocytic, edema, fibrin, E, M3  Keratitis – ulcerative, granulocytic, LP, edema, E, M3  Anterior uveitis – granulocytic, fibrin, D, M2  Dacryoadenitis – granulocytic, necrotising, hemorrhage, fibrin, D, M2  Periocular cellulitis - granulocytic, LP, edema, E, M2 | Splenitis –granulocytic, necrotising with perivascular edema, MF, M1 | Interstitial nephritis – granulocytic, necrotising, fibrin, MFE, M3  Renal tubular necrosis – MF, M2 | NA | NVL | Olfactory neuritis and perineuritis – granulocytic and necrotising, E, M2  Meningeal  lipofuscinosis – MF, M1 | Sinusitis – granulocytic, LP, necrotising, edema, MF, M2 | Hepatic necrosis – F, M1 | Stomatitis –granulocytic, fibrino-necrotising FE, M3 | Testicular spermatogenesis |
| 10446.16 | 221 | 1176 | Female | Adult | 27-2-2015 | Euthanased | Unilateral swollen eyelid and cornea  Tan plaques -skin ventral hind legs  Hydropericardium M3  Pleural, bladder, renal capsular, large intestinal edema M3 | Blepharitis and conjunctivitis – ulcerative, LP, edema, hemorrhage,  MF, M2  Anterior uveitis – LP, fibrin, edema, F, M1  Dacryoadenitis – LP, necrotising, D, M1-2 | Splenitis – granulocytic, fibrinonecrotising, F, M1-2 | Interstitial nephritis – granulocytic, necrotising, MFE, M3  Renal tubular loss and dilation | Cystitis – ulcerative, granulocytic, necrotising, edema, D, M2-3 | Intravascular thrombi – epicardial, aortic adventitia, myocardium | Meningeal and cerebral hemosiderosis – MF, M1  Optic nerve perivascular macrophages with lipofuscin and iron – MR, M1 | Sinusitis – granulocytic, LP, necrotising, FE, M2  Pleural edema – D, M2  Broncho-interstitial pneumonia – granulocytic, necrotising, fibrin, F, M1 | NVL | Stomatitis – granulocytic, LP, necrotising, F, M1  Intestinal mural edema – FE, M2 | Oophoritis – granulocytic, necrotising, -  ovum into coelomic cavity, E, M2  Epidermal necrosis – ulcerative, fibrin, MF M3 (ventral thigh)  Protozoa in urinary bladder and renal tubular lumens  Epicarditis – M3, MFE  Myocardial hydropic degeneration – hydropic, MFE  Coelomitis – granulocytic, fibrino-necrotising, MF, M2  Myodegeneration – MF, M1 |
| 10446.17 | 193 | 612 | Male | Adult | 27-2-2015 | Euthanased | Bilateral eyelid swelling – M2  Tan lesions tail and tongue  Renal parenchyma is pale  White foci on spleen | Blepharitis and conjunctivitis – granulocytic, necrotising, edema, fibrin, E, M3  Keratitis – ulcerative, granulocytic, LP, edema, E, M3  Anterior uveitis – granulocytic, LP, E, M2  Dacryoadenitis – granulocytic, fibrinonecrotising, D, M3  Periocular cellulitis - granulocytic, LP, edema, E, M2 – with osteolysis of scleral ossicles – F, M1 | Splenitis - fibrinonecrotising and granulocytic, MFE, M2 | Interstitial nephritis – granulocytic, necrotising, MF, M2  Renal tubular necrosis. Collecting ducts not evident | Cystitis – granulocytic, D, M1 | Intravascular thrombi: liver | Meningoencephalitis – LP, FE, M2  Encephalomalacia – S, M2  Meningeal lipofuscinosis – MF, M2 | Sinusitis – ulcerative, granulocytic, necrotising, ME, M2  Interstitial edema – D, M1 | Liver – thrombi MF, M2 | Stomatitis – ulcerative, granulocytic, necrotising, edema, MF, M1  Gastroenteritis – granulocytic, fibrinonecrotising, MF, M1 | Testicular spermatogenesis  Myodegeneration – MF, M1  Myocardial degeneration – MF, M1  Epidermal ulceration– granulocytic – MF, M2 (tail)  Coelomitis – MF, M1 |
| 10446.18 | 154 | 306 | Male | Adult | 27-2-2015 | Euthanased | Bilateral eyelid swelling – M2  Large spleen | Blepharitis and conjunctivitis – granulocytic, LP, necrotising, edema, E, M1  Keratitis – granulocytic, LP, edema, E, M1  Dacryoadenitis – LP, macrophages, necrotising, D, M3  Anterior chamber and lens not represented | Splenitis – granulocytic, edema, MF, M1  Well populated with lymphoid tissue | Tubulointerstitial nephritis – granulocytic, LP, necrosis, MF, M1 | Cystitis – granulocytic, LP, edema, D, M1 | Intravascular thrombi: dermal, serosal  Vasculitis: liver | Hydrocephalus and thin cortex.  Meningeal heterophils scattered ventral meninges. | Sinusitis – non-suppurative to fibrino-necrotising, necrotising – E, M2  Lung - NA | Hepatitis – granulomatous, MF, M1 | Stomatitis – LP, necrotising, E, M2  Gastroenteritis – granulocytic, LP, D, M1-2  Duodenitis – granulocytic, necrotising, serosal thrombi – D, M2 | Epidermal oedema – MF, M1  Myodegeneration – MF, M1  Coelomitis – MF, M1 |
| 10446.19 | 228 | 1080 | Female | Adult | 27-2-2015 | Euthanased | Bilateral eyelid swelling – M2  Hydropericardium | Blepharitis and conjunctivitis – ulcerative, granulocytic, fibrinonecrotising, E, M3  Corneal necrosis – F, M2  Dacryoadenitis – granulocytic, LP, fibrinonecrotising, D, M3  Uveitis – LP, MF, M1 | Splenitis – granulocytic and fibrinonecrotising, edema, MFE, M2 | Interstitial nephritis – granulocytic , necrosis, edema, fibrin – MFE, M2  Large collecting ducts not visible. Small ducts with dysplastic epithelium | Cystic mural edema – D, M2 | Vasculitis – fibrinoid with endothelial proliferation, MF, M1  Possible early thrombotic change: liver, lung, intestinal serosa | Meningeal and cerebral lipofuscinosis – E, M2 | Sinusitis – LP, necrotising, hemorrhagic, E, M2, | Hepatitis – granulocytic, fibrinonecrotising, F, M1 | Stomatitis –LP, edema, MF, M1  Intestinal vasculitis- granulocytic, necrotising, MF, M1 | Myocardial degeneration – S, M2  Myodegeneration and myositis – granulocytic, necrotising, E, M2 |
| 10446.20 | 220 | 955 | Female | Adult | 27-2-2015 | Euthanased | Bilateral eyelid swelling – M2  Tan plaques legs  White foci spleen and kidneys  Mesenteric edema | Blepharitis and conjunctivitis – ulcerative, granulocytic, fibrinonecrotising, E, M3  Uveitis and choroiditis – granulocytic, fibrinonecrotising, hemorrhage, E, M3  Keratitis – ulcerative, granulocytic, FE, M2  Dacryoadenitis – granulocytic, fibrinonecrotising, hemorrhage, E, M3  Intra-ocular haemorrhage – E, M2 | Splenitis – granulocytic, fibrinonecrotising, MF,M2 | Tubulo-interstitial nephritis – granulocytic, fibrinonecrotising, MF, M1  Dysplastic to necrotic collecting duct epithelium – MF, M2 | NA | Intravascular thrombi: spleen, trachea | Meningeal haemorrhage – MF, M2  Meningeal lipofuscinosis – MF, M1 | Sinusitis – granulocytic, LP, necrotising, MF, M2  Tracheitis – granulocytic, necrotising, F, M1  Tracheal adventitial thrombi MF, M1  Lung NA | Hepatitis – fibrinonecrotising and granulocytic, F, M1 | Stomatitis – granulocytic, LP, fibrinonecrotising, E, M2  Colitis – granulocytic, MF, M1 | Oophoritis – fibrinonecrotising and granulocytic, S, M2  Epidermal necrosis – fibrino-ulcerative – granulocytic, MF, M3  Dermatitis and epidermitis – granulocytic, MF, M2  Myocardial degeneration – MF, M1  Myodegeneration – MF, M1  Myositis- interstitial, granulocytic, F, M1 (Thigh) |
| 14617.1  2018 release | 142 | 253 | Male | Subadult | 4-5-2022 | Euthanased  Scratching right eye | Bilateral eyelid swelling and tan exudate  Blocked nostril  Mesenteric edema | Blepharitis and conjunctivitis –granulocytic, fibrin, edema, E, M2  Dacryoadenitis – granulocytic, necrotising, D, M3  Cornea and uvea not represented | NVL | Nephritis, granulocytic, necrotising, MF, M2  Embolic pattern | NA | Embolic pattern of lung, renal and hepatic lesions | Optic neuritis and encephalitis – LP, granulocytic, MF, M1 | Sinusitis – granulocytic, necrotising, MFE, M3  Broncho-interstitial pneumonia - granulocytic, necrotising, MF, M2 | Portal hepatitis - granulocytic, LP, MF, M1  Hepatocellular degeneration - MF, M1 | Esophagitis – granulocytic, necrotising, MFE, M2  Enteritis – granulocytic, LP, necrotising, MF, M2 | Oral and gastric mural granulomas – F, M1 - insect matter |
| 14617.2  2018 release | 137 | 255 | Male | Subadult | 4-5-2022 | Euthanased | Bilateral eyelid swelling and tan exudate,  Miliary tan foci dorsal tongue, liver, spleen, intestinal serosa  Mesenteric edema | Blepharitis and conjunctivitis –epidermal hyperkeratosis, edema, granulocytic exudate, D, M2  Keratitis – ulcerative, necrotising, granulocytic, edema, FE, M2  Anterior uveitis – granulocytic, fibrin – FE, M2  Dacryoadenitis– granulocytic, necrotising – D, M3 | NVL | Nephritis – granulocytic, necrotising, MF, M2  Embolic pattern | NA | Embolic pattern of renal and hepatic lesions | Meningo- encephalitis granulocytic MF, M1 | Sinusitis – granulocytic, necrotising, MFE, M3  Tracheitis – segmental, granulocytic, necrotising, MFE, M2-3  Broncho-interstitial pneumonia, granulocytic, necrotising, MF, M2  Pleural edema – D, M1 | Hepatitis – granulocytic, necrotising, MFE, M2 | Stomatitis – granulocytic, necrotising, MF, M1  Esophagitis, gastritis, and enteritis – granulocytic, necrotising, MF, M2  Enteritis – granulocytic, LP, necrotising, MF, M2-3 | Thymitis – granulocytic, F, M1  Dermatitis and epidermitis – granulocytic, F, M1 |
| 14617.3  2018 release | 128 | 198 | Male | Subadult | 4-5-2022 | Euthanased | Bilateral eyelid swelling with ulcerated lower lids and exudate  Mesenteric edema | Blepharitis and conjunctivitis – ulcerative, granulocytic, fibrinonecrotising, D, M3  Keratitis – granulocytic, edema, F, M1  Uveitis - granulocytic, F, M1  Dacryoadenitis– granulocytic, necrotising – D, M3 | NVL | Nephritis – granulocytic, necrotising, MF, M2  Embolic pattern | NA | Embolic pattern of renal and hepatic lesions |  | Sinusitis – granulocytic, necrotising, MFE, M3  Broncho-interstitial pneumonia – granulocytic, necrotising, MF, M2 | Hepatitis – granulocytic, necrotising, MF, M2 | Stomatitis – granulocytic, necrotising, MF, M1-2  Esophagitis, gastritis, and enteritis – granulocytic, necrotising, MF, M2-3  Mesenteric and serosal oedema – MFE, M2 | Periocular dermatitis – F, M3 |
| 14617.4  2018 release | 118 | 150 | Male | Subadult | 6-5-26 | Euthanased | Bilateral eyelid swelling and tan exudate,  Mesenteric edema  Small green focus right lung cranial  Focal, full thickness skin ulcers – crusted and contracted – right neck and thigh (wounds) | Blepharitis and conjunctivitis - granulocytic, edema, MF, M2  Granulocytic exudate in conjunctival sulcus  Keratitis, uveitis - granulocytic, MFE, M2  Dacryoadenitis– granulocytic necrotising – D, M2-3 | NVL | Nephritis – granulocytic, necrotising, MF, M2 | NA | Embolic pattern of renal and hepatic lesions | Neuritis, encephalitis - granulocytic, necrotising, MF, M2-3 | Sinusitis – granulocytic, necrotising, MFE, M2-3  Tracheitis and broncho-interstitial pneumonia – granulocytic, necrotising, MF, M2 | Hepatitis – granulocytic, MF, M1 | Stomatitis (deep lamina propria) - granulocytic, necrotising, F, M2-3  Esophagitis, gastritis, and enteritis – granulocytic, LP, necrotising, MF, M2-3 |  |
| 14617.5  2018 release | 138 | 237 | Male | Subadult | 6-5-26 | Euthanased | Bilateral eyelid swelling and tan exudate,  Focal tan skin lesions tail and ventral hindlimbs  Miliary tan foci dorsal tongue, liver, lungs | Blepharitis and conjunctivitis - granulocytic, edema, MF, M2  Granulocytic exudate in conjunctival sulcus  Keratitis, uveitis - granulocytic, MF, M1  Dacryoadenitis– granulocytic necrotising – D, M2-3 | NVL | Nephritis – granulocytic, necrotising, MF, M2 | NA | Embolic pattern of renal and hepatic lesions | Encephalitis – granulocytic, MF, M1-2 | Sinusitis – granulocytic, necrotising, MFE, M2  Tracheitis– granulocytic, necrotising, MFE, M2  Broncho-interstitial pneumonia – granulocytic, necrotising, F, M2 | Hepatitis – granulocytic, MF, M2 | Stomatitis - granulocytic, necrotising, F, M2-3  Esophagitis, gastritis, and enteritis – granulocytic and necrotising, MF, M2-3  Mesenteric oedema – S, M2 | Dermatitis and epidermitis – granulocytic, F, M1 (Thighs) |
| 15811.2 | 150 | 247 | Female | Subadult | 25-4-2024 | Euthanased | Bilateral chemosis m3, unilateral erythema  Pulmonary consolidation  Splenomegaly  Mesenteric, pleural oedema  Hydropericardium | Blepharitis and conjunctivitis – granulocytic, fibrin, edema, D, M3  Keratitis, scleritis - granulocytic, F, M1  Dacryoadenitis– granulocytic necrotising, D, M2-3  Periocular cellulitis - granulocytic, LP, fibrin, edema, D, M3 | NVL | Interstitial nephritis, LP, macrophages, granulocytes, MF, M2 | NA | NVL | Olfactory perineuritis – granulocytic, F, M2 | Sinusitis –granulocytic, fibrinoid, LP, MFE, M2 | NVL | Stomatitis – granulocytic, LP, F, M1  Enteritis – granulocytic, LP, F, M2  Mesenteric edema - E, M2 | Ovary inactive  Renal tubular protozoa – MF, M1 – presumed myxozoa  Epicardial haemorrhage – F, M2 |
| 15811.3 | 133 | 224 | Male | Subadult | 25-4-2024 | Euthanased | Bilateral chemosis m2,  Pulmonary consolidation  Splenomegaly  Mesenteric, pleural oedema M2  Hydropericardium | Blepharitis and conjunctivitis - ulcerative, fibrinous, granulocytic and LP, E, M3  Dacryoadenitis – granulocytic, fibrinonecrotising, D, M3  Scleritis- granulocytic, LP, F, M1  Periocular cellulitis - granulocytic, LP, edema – D, M3 | NVL | Interstitial nephritis – granulocytic, LP, edema, MF, M1 | Cystitis, granulocytic, edema, LP, D, M2 | NVL | Meningoencephalitis (olfactory lobe) – granulocytic, MF, M1 | Sinusitis – necrotising and granulocytic, LP, MF, M2 | NVL | Gastric mural edema, MF, M2  Enteritis –granulocytic, LP, edema, D, M2  Mesenteric edema – D, M2-3 |  |
| 15811.4 | 134 | 222 | Female | Subadult | 25-4-2024 | Euthanased | Bilateral chemosis m3,  Pulmonary consolidation  Splenomegaly  Mesenteric, pleural oedema  Hydropericardium | Blepharitis and conjunctivitis – granulocytic, LP fibrinous, E,M3  Dacryoadenitis – granulocytic, LP, fibrinonecrotising, epithelial dysplasia, D, M3  Periocular cellulitis - granulocytic, LP, edema, E, M3 | NVL | Interstitial nephritis, LP, MF, M1 | Cystitis - granulocytic, LP, D, M1 | Hepatic vasculitis – MF,M2 | NVL | Sinusitis – granulocytic LP, MF, M1  Interstitial pneumonia – granulocytic, LP , F, M1 | Hepatic hemorrhage and LP vasculitis – MF, M2 | Gastritis - granulocytic, edema, MF, M1-2  Mesenteric edema – D, M2-3 | Ovary - inactive  Renal tubular protozoa – MF, M1 – presumed myxozoa  Coelomitis – LP, FE, M1 |
| 15811.5 | 144 | 274 | Male | Subadult | 25-4-2024 | Euthanased | Bilateral chemosis m3,  Pulmonary consolidation  Splenomegaly  Mesenteric, pleural oedema  Hydropericardium | Blepharitis – granulocytic, LP, edema, E, M1-2  Dacryoadenitis – necrotising, granulocytic, LP, D, M3  Corneal ulceration, scleral erosion, F, M1-2  Periocular cellulitis - granulocytic, LP, edema, E, M3 | NVL | NVL | NA | Vasculitis - hepatic, F, M1 | Olfactory perineuritis – granulocytic MF, M1 | Sinusitis – granulocytic LP, MF, M2 | Hepatocellular degeneration and necrosis – granulocytic LP, MF, M1 | Gastritis - granulocytic, edema, MF, M1-2  Mesenteric edema – D, M2-3 | Testicle – no spermatogenesis  Duodenal coccidiosis – F, M1 |
| 15811.6 | 130 | 203 | Female | Subadult | 25-4-2024 | Euthanased | Bilateral chemosis  Pulmonary consolidation  Splenomegaly  Mesenteric, pleural oedema  Hydropericardium | Blepharitis and conjunctivitis – ulcerative, granulocytic, LP, edema, fibrin – MF, M2  Dacryoadenitis – necrotising, granulocytic, D, M2  Periocular cellulitis - granulocytic, LP, edema, E, M2 | NVL | Interstitial nephritis – granulocytic – radiating hilus to cortex – MF, M1 | Cystitis – granulocytic, LP, D, M1-2 | Pulmonary vasculitis | NVL | Sinusitis – LP, MF, M1  Interstitial pneumonia, granulocytic, F, M1 | Sinusoidal clusters of granulocytes, LP, MF, M1 | Enteritis and colitis – granulocytic, LP, D, M2  Mesenteric and serosal edema, D, M1 | Ovary inactive  Renal tubular protozoa – MF, M1 – presumed myxozoa  Granuloma in sinus lamina propria, F, M1  Gastric mural granuloma – F, M1 - insect matter |
| 15811.7 | 135 | 202 | Female | Subadult | 25-4-2024 | Euthanased | Bilateral chemosis  Pulmonary consolidation  Splenomegaly,  Mesenteric, pleural oedema  Hydropericardium | Blepharitis and conjunctivitis – ulcerative, granulocytic, LP, edema, fibrin – MF, M1-2  Dacryoadenitis – necrotising, granulocytic, D, M2  Periocular cellulitis - granulocytic, LP edema, E, M2 | NVL | Interstitial nephritis – granulocytic – radiating hilus to cortex – MF, M1 | Cystitis – granulocytic, D, M! | NVL | NVL | Sinusitis – granulocytic, edema, LP, MF, M1 | Sinusoidal clusters of granulocytes, LP, MF, M1 | Dilated intestinal lacteals  Enteric granulocytic, LP infiltrates, MFE, M1 | Ovary inactive  Perioral, gastric mural, splenic, mesenteric granulomas – F, M1 - insect matter  Renal tubular protozoa – MF, M1 – presumed myxozoa |
| 15811.8 | 132 | 194 | Female | Subadult | 25-4-2024 | Euthanased | Bilateral chemosis and erythema  Pulmonary consolidation  Splenomegaly,  Mesenteric, pleural oedema  Hydropericardium | Blepharitis and conjunctivitis – granulocytic, LP, edema, fibrin, MF, M2  Dacryoadenitis – necrotising, granulocytic, D, M2  Periocular cellulitis - granulocytic, LP, edema, E, M2 | NVL | Interstitial nephritis – LP, MF, M1 | Cystic mucosal ulceration, S, M2  Cystic mural edema - D, M3 | NVL | NVL | Tracheitis – granulocytic, F, M1 | NVL | Enteritis – granulocytic, LP, D, M2  Mesenteric and serosal edema, D, M1 | Ovary inactive  Renal tubular protozoa, MF, M1 – presumed myxozoa  Perioral granulomas, F, M2 – insect matter |
| 15811.9 | 148 | 276 | Unknown | Subadult | 25-4-2024 | Euthanased | Bilateral chemosis and erythema  Pulmonary consolidation  Splenomegaly  Mesenteric, pleural oedema  Hydropericardium | Blepharitis and conjunctivitis – ulcerative, granulocytic, LP, edema, fibrin, MF, M2  Dacryoadenitis – necrotising, granulocytic, D, M2-3  Periocular cellulitis - granulocytic, LP, edema, E, M2 | NVL | Interstitial nephritis – LP, MF, M2 | Cystic mucosal ulceration -S, M2  Cystic mural edema, D, M3 | Vasculitis – perineural, hepatic – granulocytic, MF, M1 | NVL | Tracheitis – granulocytic, F, M2 | Hepatic vasculitis - granulocytic, MF, M1 | Mesenteric and serosal edema - F, M1 | Renal tubular protozoa - MF, M1 – presumed myxozoa  Perioral granulomas -F, M2 – insect matter |
| 15811.10 | 150 | 293 | Female | Subadult | 25-4-2024 | Euthanased | Bilateral chemosis and erythema  Pulmonary consolidation  Splenomegaly  Mesenteric, pleural oedema  Hydropericardium | Blepharitis and conjunctivitis – ulcerative, granulocytic, LP, edema, fibrin, E, M2  Dacryoadenitis – granulocytic, necrotising, D, M2-3  Periocular cellulitis - granulocytic, LP, edema, E, M2 | NVL | Interstitial nephritis – granulocytic, MF, M1  Collecting duct epithelial necrosis – MF, M1 | Cystic mural edema, D, M3 | Vasculitis – periesophageal – fibrinous, F, M2 | NVL | Sinusitis – granulocytic, LP, MF, M1 | NVL | Peri-esophageal vasculitis -fibrinous, F, M2  Gastritis – granulocytic, MF, M2  Enteritis – LP, S, Mw  Mesenteric edema - MF, M1 | Ovary inactive  Perioral granulomas, urinary bladder -F, M2 – insect matter  Duodenal coccidiosis – MF, M1 |
| 15811.11 | 139 | 220 | Male | Subadult | 25-4-2024 | Euthanased | Bilateral chemosis  Pulmonary consolidation  Splenomegaly  Mesenteric, pleural oedema  Hydropericardium | Blepharitis and conjunctivitis – granulocytic, LP, edema, fibrin, E, M2  Dacryoadenitis – granulocytic, LP, D, M2-3  Periocular cellulitis - granulocytic, LP, edema, E, M2 | NVL | Interstitial nephritis – LP, MF, M1 | Cystic mural edema, D, M3 | NVL | Olfactory nerve perineuritis – LP, granulocytic, F, M1 | Sinusitis – granulocytic, LP, MF, M1 | NVL | Mesenteric edema - MF, M1 | Testicle – no spermatogenesis  Renal tubular protozoa - MF, M1 – presumed myxozoa  Duodenal coccidiosis – F, M1 |
| 15811.12 | 139 | 240 | Female | Subadult | 25-4-2024 | Euthanased | Bilateral chemosis m 3, tan exudate  Pulmonary consolidation D, M2  Splenomegaly – M2,  Mesenteric, pleural oedema M2  Hydropericardium – M1 | Blepharitis and conjunctivitis – granulocytic, LP, edema, fibrin, E, M1-2  Dacryoadenitis – granulocytic, LP, fibrin, D, M2-3  Periocular cellulitis - LP, granulocytic, LP, edema, E, M2 | NVL | Granulocytes scattered in collecting duct epithelium | Cystitis – mural oedema, granulocytes, LP, M2 | NVL | NVL | Sinusitis – granulocytic, LP, MF, M1  Tracheitis – granulocytic, S, M1 | Biliary duct single cell necrosis – MF, M1 | NVL | Ovary inactive  Perioral and gastric granulomas - F, M2 – insect matter  Duodenal coccidiosis – F, M1 |
| 15811.13 | 120 | 144 | Male | Subadult | 25-4-2024 | Euthanased | Bilateral chemosis with tan exudate  Pulmonary consolidation  Splenomegaly  Mesenteric, pleural oedema  Hydropericardium | Blepharitis and conjunctivitis – granulocytic, LP, edema, fibrin, E, M2  Dacryoadenitis – granulocytic, LP, fibrinonecrotising, D, M2-3  Periocular cellulitis - granulocytic, LP, edema, E, M2 | NVL | Interstitial nephritis – LP, MF, M1  Periglomerular fibrosis – MF, M1 | Cystic mural edema – D, M3  Cystitis –granulocytic, MF, M1 | NVL | NVL | Sinusitis – granulocytic, LP, MF, M1 | NVL | Stomatitis – granulocytic, F, M1  Esophagitis – LP, MF, M1  Mesenteric edema – S, M1 | Testicle – no spermatogenesis  Orchitis – granulocytic, MF, M1  Perioral and gastric granulomas - F, M2 – insect matter  Renal tubular protozoa - MF, M1 – presumed myxozoa  Splenic capsular granuloma – F, M1 |
| 15811.15 | 161 | 385 | Male | Adult | 30-4-2024 | Found Dead | Bilateral chemosis  Broncho-pneumonia – exudative  Peri-hepatitis - fibrinous,  Splenomegaly  Mesenteric oedema | Blepharitis and conjunctivitis – granulocytic, LP, edema, fibrin, E, M2  Keratitis – granulocytic, MF, M1  Dacryoadenitis – granulocytic, LP, fibrinonecrotising, D, M2-3  Periocular cellulitis - LP granulocytic, edema, osteolysis, E, M2 | Splenitis - granulocytic, necrotising, MF, M2 | Interstitial nephritis – LP, MF, M1 | Cystitis – granulocytic, D, M1 | NVL | NVL | Tracheitis – necrotising, granulocytic, MF, M2  Bronchitis – necrotising, granulocytic, multinucleated giant cells, D, M2-3 | Hepatocellular atrophy - D, M1 | Moderate autolysis | Testicular spermatogenesis  Coelomitis – granulocytic, MF, M2 |
| 15811.17 | 144 | 259 | Male | Adult | 27-4-2024 | Treated and Died  Frozen-thawed remains  Treated 1-2 days | Bilateral chemosis M2  Duodenal haemorrhage  Splenomegaly  Mesenteric oedema | Blepharitis and conjunctivitis – ulcerative, necrotising, granulocytic, edema, fibrin, E, M2  Dacryoadenitis – necrotising, D, M2-3 | NVL | NVL | NA | NVL | NVL | Sinus hemorrhage  Tracheitis – granulocytic, MF, M2  Interstitial pneumonia – granulocytic, F, M1 | NVL | Moderate autolysis  Mesenteric edema – MF, M2 | Testicular spermatogenesis  Duodenal coccidiosis |
| 15811.18 | 185 | 670 | Female | Subadult | 14-5-2024 | Treated 18 days then Died  Frozen-thawed remains  Treated 18 days.  Mild eye lesions that resolved.  Systemic swelling | Bilateral chemosis M2  Pulmonary consolidation  Hydropericardium  Tan splenic foci  Renal pallor  Pleural and mesenteric oedema | Blepharitis and conjunctivitis –edema, E, M2  Dacryoadenitis – granulocytic, necrotising, D, M3  Periocular cellulitis - granulocytic, edema, MF, M2 | Splenitis – granulocytic, necrotising, MF, M3 | Protein losing nephropathy – MF, M2  Urate nephrosis – MF, M2 | NVL | NVL | NVL | Sinusitis – granulocytic, necrotising, MF, M2  Tracheitis - granulocytic, necrotising, D, M3  Interstitial pneumonia – granulocytic, MF, M3 | Hepatic necrosis – granulocytic, MF, M2 | Moderate autolysis  Stomatitis – granulocytic, necrotising, F, M2 | Ovary inactive  Renal tubular protozoa - MF, M1 – presumed myxozoa  Sinus metazoan parasite  Pancreatic necrosis – granulocytic, MF, M2 |
| 15811.19 | 180 | 457 | Female | Subadult | 20-5-2024 | Treated 26 days then died.  Frozen-thawed remains | Chemosis  Keratitis  Stomatitis  Splenic and renal pallor  Perihepatic fibrosis | Blepharitis and conjunctivitis – ulcerative, necrotising, granulocytic, edema, fibrin, D, M2  Keratitis – granulocytic, fibrinous, D, M3  Dacryoadenitis – necrotising, D, M2-3 | Splenic capsular necrosis, granulocytes – D, M2 | Protein losing nephropathy – MF, M2 | NA | Vasculitis – mesenteric, gastric, bone marrow, skeletal muscle, peri-oral, peri-ocular, - granulocytic, fibrinonecrotising, MFE, M3 | Meningoencephalitis – granulocytic, E, M3 | Broncho-interstitial pneumonia – granulocytic, MF, M3 | Hepatic necrosis – granulocytic, MF, M2 | Moderate autolysis  Gastric mural vasculitis with partial thrombi  Gastrointestinal luminal acid haematin – E, M2 | Ovary inactive  Coelomitis – splenic capsule – necrotising, granulocytic, E, M3  Renal tubular protozoa - MF, M1 – presumed myxozoa  Thymitis – granulocytic, MF, M2 |
| 15811.20 | 263 | 141 | Male | Subadult | 10-5-2024 | Treated 10 days then euthanased  Frozen-thawed remains  Purulent nasal discharge cleared. | Chemosis  Keratitis – D, M3  Stomatitis  Splenic and renal pallor  Tan foci epicardium and renal interstitium | Blepharitis and conjunctivitis – granulocytic, ulcerative, necrotising, edema, fibrin, D, M3  Keratitis – granulocytic, fibrinous, D, M3, fungal invasion  Anterior uveitis – granulocytic, necrotising, D, M3  Dacryoadenitis – granulocytic, necrotising, D, M3  Periocular cellulitis - LP granulocytic, edema, osteolysis, E, M2 | Splenic fibrin deposition – F, M1 | Interstitial nephritis – granulocytic, MF, M2  Collecting duct necrosis, granulocytes, MF, M2  Urate nephrosis – MF, M1 | NA | NVL | Meningoencephalitis – granulocytic, E, M3 | Sinusitis – granulocytic, MFE, M2  Tracheitis – granulocytic, MF, M1  Interstitial pneumonia – granulocytic, F, M1 | Hepatic necrosis – granulocytic, E, M2 | Moderate to marked autolysis  Stomatitis – ulcerative, necrotising, granulocytic, D, M3  Esophagitis – granulocytic, MF, M2  Gastritis | Epicarditis and myocarditis granulocytic, MF, M2  Thymitis – granulocytic, MF, M2  Osteomyelitis and osteolysis – granulocytic, MF, M2  Renal tubular protozoa - MF, M1 – presumed myxozoa |
| 15811.21 | 158 | 210 | Male | Subadult | 17-5-2024 | Treated 16 days then died.  Frozen -  thawed remains  Mild chemosis, which resolved then recurred | Chemosis  Stomatitis  Tan foci myocardium, kidney  Splenic pallor | Blepharitis and conjunctivitis – ulcerative, necrotising, granulocytic, edema, fibrin, D, M3  Keratitis – granulocytic, fibrinous, D, M3 –fungal invasion  Anterior uveitis – necrotising, hemorrhagic, granulocytic, D, M3  Dacryoadenitis – necrotising, D, M3  Periocular cellulitis - LP granulocytic, edema, osteolysis, E, M2 | Splenic capsular necrosis, granulocytes – S, M2 | Interstitial nephritis – granulocytic, MF, M2  Urate nephrosis – MF, M1 | NA | Vasculitis – oral, MF, M2-3 | Meningoencephalitis – granulocytic, E, M3 | Sinusitis – granulocytic, E, M3  Tracheitis and interstitial pneumonia – granulocytic, E, M3 | Hepatic necrosis – granulocytic, E, M2 | Moderate to marked autolysis  Stomatitis – ulcerative, necrotising, granulocytic, D, M3, fungal and bacterial invasion  Gastritis – granulocytic, edema, D, M2 | Orchitis – granulocytic, LP, necrotising – D, M3  Myocarditis – granulocytic, MF, M2-3  Thymitis – granulocytic, MF, M2  Coelomitis – necrotising, fibrinous, E, M3  granulocytic, necrotising, MF, M2-3  Perioral granuloma – F, M1 – insect matter |
| 15811.24 | 115 | 132 | Unknown | Juvenile |  | Treated then died  Frozen -  thawed remains | Chemosis – D, M3  Stomatitis – D, M3 | Blepharitis and conjunctivitis – ulcerative, necrotising, granulocytic, edema, fibrin, D, M3  Dacryoadenitis – granulocytic, necrotising, D, M3  Periocular cellulitis - LP granulocytic, necrotising, edema, fibrin, E, M2 | Splenitis– granulocytic, necrotising, MF, M2 | Interstitial nephritis -granulocytic, MF, M2  Protein losing nephritis – MF, M1 | NA | Vasculitis – oral, | Encephalitis – granulocytic, necrotising, MF, M2 | Sinusitis – granulocytic, E, M3  Tracheitis – granulocytic, mucosal sloughing into airways, D, M3 | Hepatitis– granulocytic, necrotising, MF, M2 | Moderate to marked autolysis  Stomatitis – ulcerative, , granulocytic, necrotising, D, M3, fungal and bacterial invasion  Gastritis – granulocytic, S, M2 | Perioral granuloma – F, M1 – insect matter |
| 15811.25 | 105 | 79 | Male | Juvenile |  | Treated then died  Frozen -  thawed remains | Emaciation  Dehydration  Sunken eyelids (examination not possible) | Blepharitis and conjunctivitis – ulcerative, necrotising, granulocytic, edema, fibrin, D, M3  Dacryoadenitis –granulocytic, necrotising, D, M3  Periocular cellulitis - LP granulocytic, necrotising, edema, E, M2 | Splenitis – granulocytic, necrotising, MF, M2 | NVL | NA | Vasculitis | NVL | Sinusitis – granulocytic, fibrinonecrotising, MF, M2 | NVL | Stomatitis – granulocytic, necrotising, D, M3  Mesenteric and serosal edema – D, M2 | Testicle – no spermatogenesis  Perioral granuloma – F, M1 – insect matter |
| 15811.26 | 162 | 378 | Male | Adult | 26-6-2024  Treated 13 days | Treated then died  Frozen -  thawed remains | Chemosis  Stomatitis  Hydropericardium – | Blepharitis and conjunctivitis – ulcerative, granulocytic, necrotising, edema, fibrin, D, M3  Dacryoadenitis – granulocytic, necrotising, D, M3  Periocular cellulitis - LP granulocytic, necrotising, edema, E, M2 | Splenitis – granulocytic, necrotising, MF, M2-3 | Acute renal tubular necrosis – MF, M2  Protein losing nephritis – MF, M1 | NA | Vasculitis and thrombosis - perioral | Meningitis – granulocytic, fibrinous, F, M1 | Sinusitis – granulocytic, fibrinonecrotising, MF, M2  Tracheitis - granulocytic, necrotising, MF, M2-3  Interstitial pneumonia - granulocytic, necrotising, MF, M2-3 | Hepatitis – granulocytic and necrotising, MF, M2-3 | Stomatitis – thrombosis, vasculitis, necrosis, granulocytic, MF, M2 | Testicular spermatogenesis  Adrenitis - granulocytic, necrotising, MF, M2-3 |

NA – not available,

NVL – no visible lesions in the sections examined

MS – multisystemic

D – diffuse

E - extensive

MFE – multifocally extensive

M1 – mild

M2 - moderate

M3 – marked

LP – lymphoplasmacytic infiltrates
