## Supplementary Tables S4 for "Bellinger River Virus 2: The Emergence of a Second Novel Pathogenic Nidovirus in Bellinger River Snapping Turtles (*Myuchelys georgesi*)"

**Table S4.** Summary of Bellinger River snapping turtle pathology

|  | **2015**  **(n=15)** | **2022 (n=5)** | **2024**  **(n=21)** |
| --- | --- | --- | --- |
| Sex  (male/female/unknown) | 11/4/0 | 0/3/2 | 10/9/2 |
| Age Class (Adult/subadult/ juvenile) | 15/0/0 | 0/5/0 | 2/17/2 |
| Blepharitis, chemosis, periocular cellulitis* | 15 | 5 | 21 |
| Dacryoadentits^#^ | 15 | 5 | 21 |
| Keratitis/uveitis/choroiditis^#^ | 14/14/3 | 4/4/2^†^ | 6/2/0 |
| Sinusitis* | 11 | 5 | 17 |
| Stomatitis* | 14 | 5 | 9 |
| Skin lesions (excluding periocular)* | 8 | 2 | 0 |
| Hydropericardium* | 1 | 0 | 14 |
| Splenitis* | 14^†^ | 0 | 6 |
| Hepatitis* | 9 | 4 | 10 |
| Nephritis/tubular necrosis* | 13 | 4 | 15 |
| Cystitis* | 13^†^ | - | 11^††^ |
| Thrombosis/vasculitis ^#^ | 13/2 | 5/3 | 0/10 |
| Oesophagitis/gastroenteritis^#^ | 0/12 | 3/3 | 2/11 |
| Tracheitis/bronchitis/interstitial pneumonia# | 1/3/4^§^ | 5/5/5 | 10/2/7 |
| Central/peripheral nervous inflammation^#^ | 5/4 | 4/5 | 6/3 |
| Hydrocephalus | 3 | 0 | 0 |
| Internal genitalia inflammation^#^ | 6 | 1 | 1 |
| Epicarditis/coelomitis* | 3/4 | 0/2 | 1/5 |

*Gross and/or histological observations

^#^Histological observation

-Tissue not available for any animals in the cohort

^†^Tissue not available for one animal in the cohort

^§^Tissue not available for t animals in the cohort

^††^Tissue not available for nine animals in the cohort
